# Boosting Proteasome Activity: A Novel Mechanism of NMDAR Blockers Against Neurodegeneration

**DOI:** 10.1101/2024.08.20.608787

**Authors:** Fikret Sahin, Aslihan Gunel, Buse Turegun Atasoy, Ulku Guler, Bekir Salih, Isunsu Kuzu, Mehmet Taspinar, Ozgur Cinar, Selda Kahveci

## Abstract

NMDAR antagonists, such as memantine and ketamine, have shown efficacy in treating neurodegenerative diseases and major depression. The mechanism by which these drugs correct the aforementioned diseases is still unknown. Our study reveals that these antagonists significantly enhance 20S proteasome activity, crucial for degrading intrinsically disordered, oxidatively damaged, or misfolded proteins, factors pivotal in neurodegenerative diseases like Alzheimer’s and Parkinson’s. In a mouse model, ketamine administration notably altered brain synaptic protein profiles within two hours, downregulating proteins linked to neurodegenerative conditions. Furthermore, the altered proteins exhibited enrichment in terms related to plasticity and potentiation, including retrograde endocannabinoid signaling—a pivotal pathway in both short- and long-term plasticity that may elucidate the long-lasting effects of ketamine in major depression. Via the ubiquitin-independent 20S proteasome pathway (UIPS), these drugs maintain cellular protein homeostasis, crucial as proteasome activity declines with age leading to protein aggregation and disease symptoms. The elucidation of the mechanistic pathways underlying the therapeutic effects of NMDAR antagonists holds promise for developing new treatment strategies for brain diseases, including schizophrenia, Alzheimer’s, and Parkinson’s.

## Introduction

The proteasome system is the cell’s primary defense against proteotoxic stress from oxidative damage. Age-related dysfunction in this system disrupts crucial signaling pathways and is linked to various human diseases^1-4^. Regulating the proteolytic machinery to modulate intracellular protein concentration holds promise for developing treatments for aging-related diseases, including neurodegeneration, cancer, and autoimmunity^1-4^. The proteasome system includes the ubiquitin-proteasome system (UPS) and the ubiquitin-independent proteasome system (UIPS). Both systems degrade oxidatively damaged proteins^1-3^.

The UPS requires ubiquitination of proteins before transporting them to the 26S proteasome, consisting of the 19S regulatory particle (19S RP) and the 20S core particle (20S CP)^5^. The 19S RP recognizes a ubiquitinated substrate, removes ubiquitin, unfolds the substrate, and shuttles it into the 20S CP. Thus, the UPS involves multiple steps: ubiquitination, recognition, deubiquitination, unfolding, and degradation^5^.

In contrast, the UIPS bypasses the 19S structures of the 26S and primarily includes the 20S core and regulators such as PA28s. The 20S proteasome, constituting the core of the 26S system, is abundant in cells, making up about 1% of total cellular protein^6, 7^. While the 26S proteasome degrades ubiquitinated proteins, 20S-mediated proteolysis does not require ubiquitination. It directly degrades misfolded, oxidatively damaged, intrinsically disordered proteins (IDPs), and intrinsically disordered protein regions (IDPRs)^6, 7^. With IDRs predicted to constitute 41% of the eukaryotic proteome, the potential substrate pool for UIPS is vast^8^. These substrates, including β-amyloid (Alzheimer’s disease), Tau (Alzheimer’s disease), α-synuclein (Parkinson’s disease), and cell cycle-related proteins like p53, p27, and p21, are particularly relevant to neurodegenerative disorders^6-8^.

Despite the critical role of the 20S proteasome, few in vivo activators have been identified^9^. For the first time, our work shows that NMDAR antagonists enhance 20S proteasome activity independent of the traditional UPS pathway. This finding broadens our understanding of protein degradation, demonstrating that the 20S proteasome can degrade ubiquitinated proteins and intrinsically disordered, oxidatively damaged, and misfolded proteins, potentially with greater efficiency under certain conditions.

NMDAR antagonists, such as memantine, are effective in treating protein misfolding disorders, including Alzheimer’s disease, vascular dementia, and Parkinson’s disease^1-4^. Additionally, a single intravenous infusion of the NMDAR antagonist ketamine can rapidly improve major depression, with effects lasting up to two weeks^10^. Despite extensive research, the complete mechanism of action for these drugs remains elusive.

Synaptic modifications and alterations in synapse-associated proteins are implicated in various brain diseases^11-14^. Our study shows that ketamine administration significantly alters the synaptic protein profiles in mouse brains, downregulating 372 known synaptic proteins and upregulating 145. Enrichment analysis highlights several factors associated with cognitive, emotional, and motor phenotypes, as well as various neurological and psychiatric disorders^11-15^.

In summary, our results indicate that increased proteasome activity by NMDAR antagonists suggests a mechanistic route for their therapeutic effects on neurodegenerative diseases, previously not fully understood. This enhancement in proteasome activity also proposes new potential treatments for protein misfolding conditions, including cancer and autoimmunity.

## Results

### NMDAR antagonists reduce proteins characterized as having intrinsically disordered regions

In our laboratory studies, we found that NMDAR antagonists, ketamine and memantine, decreased levels of proteins with intrinsically disordered regions, namely p21, p27, and p53, in the cell lines tested, which included HepG2, SH-SY5Y, T98G, and Hep3B (Fig. 1A, B, Fig. S1A-C). The decrease is concentration- and time-dependent, with effects visible at 10 minutes, persisting for 6 to 8 hours, and then returning to normal levels in 12 to 24 hours, as verified by Western blot analysis (Fig. S1D-F). The concentrations of memantine (10 µM) and ketamine (500 µM) used in our experiments did not affect cell proliferation, apoptosis, or cytotoxicity, consistent with previous studies ^15, 16^. Importantly, these reductions occur independently of cycloheximide, suggesting they are not due to inhibited protein synthesis (Fig. S1G, H).

**Fig. 1.**
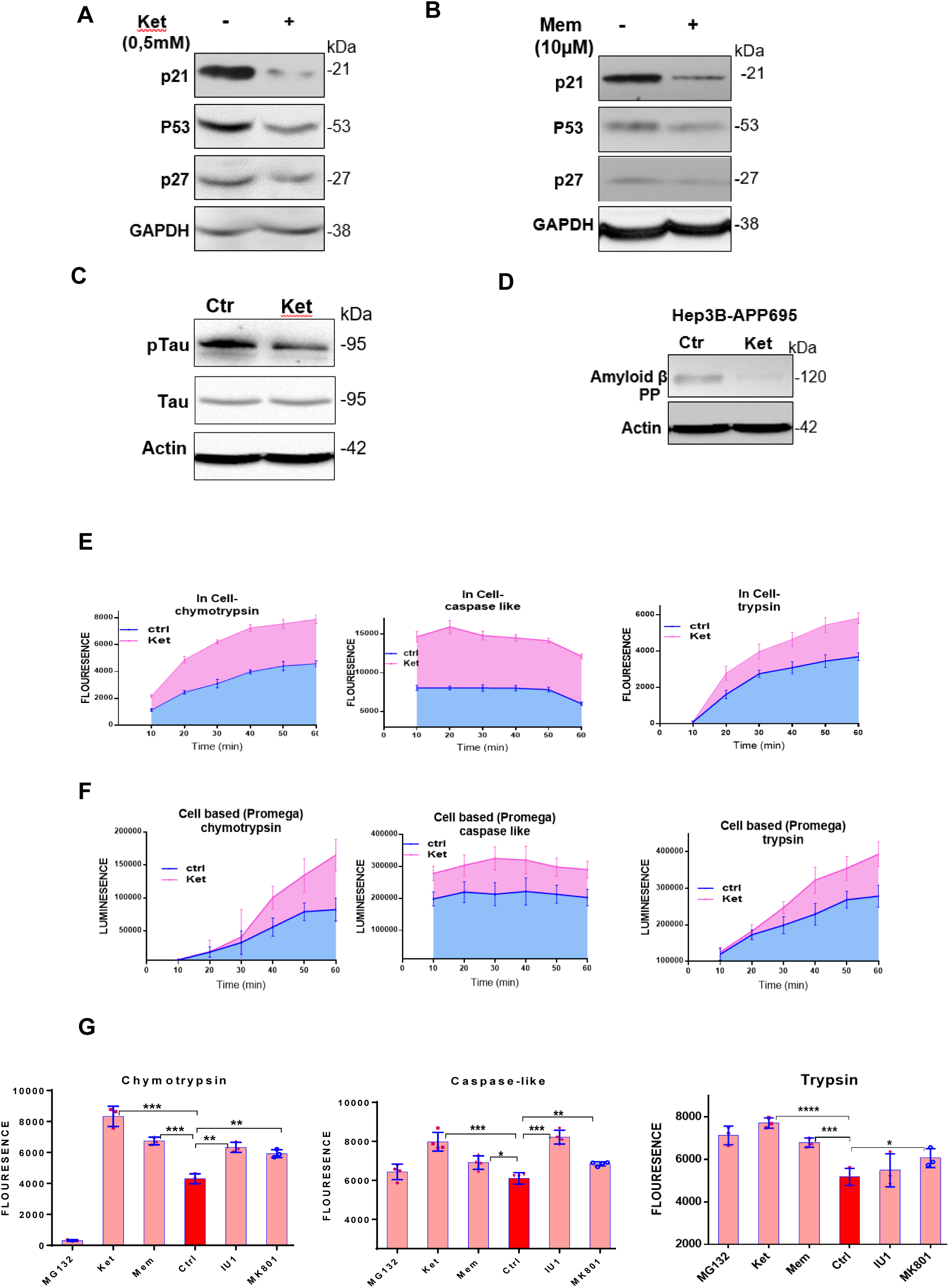
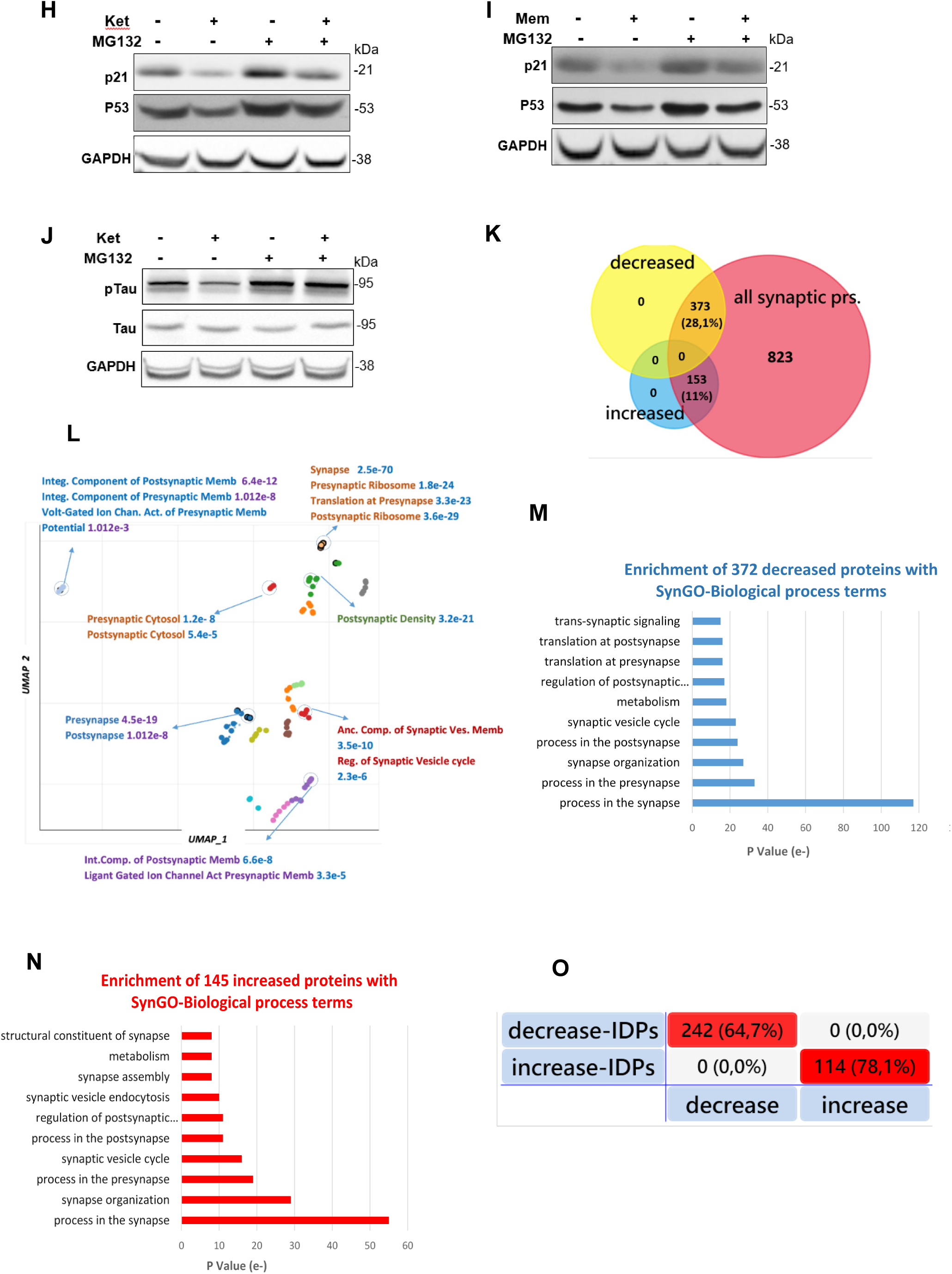
Modulation of intrinsically disordered proteins (IDPs) Levels and Proteasome Activity by NMDA Receptor Antagonists. (A) Ketamine (500µM) and (B) memantine (10µM) reduce the levels of the cell cycle IDPs p21, p53, and p27 in HepG2 cells. (C, D) Western blot analysis confirms that ketamine induces the downregulation of Alzheimer’s disease-associated intrinsically disordered proteins (IDPs), including phosphorylated Tau (pTau) in HepG2 and SH-SY5Y cells and β-amyloid precursor protein ectopically expressed in Hep3B and SH-SY5Y cells. (E) The "Intracellular Fluorogenic Proteasome Test" indicates an increase in chymotrypsin-like, caspase-like, and trypsin-like proteasome activities upon treatment with NMDA receptor antagonists. (F) Chymotrypsin-like, caspase-like, and trypsin-like proteasome activities measured in whole cells using the "Cell-Based Proteasome Luciferase System" from Promega. (G) Assessment of proteasome activity in HepG2 cells treated with additional NMDA receptor antagonists (memantine, MK801-20µM), a proteasome activator (IU1-50µM), and an inhibitor (MG132-0.5µM). (H-J) Proteasome inhibition by MG132 prevents the reduction of IDPs such as p21, p53, and pTau caused by NMDA receptor antagonists. (K-O) Protein changes in the synapse following single dose of ketamine administration (30 mg/kg) were evaluated by comparing two groups, each composed of three mice. To identify the protein alterations attributed to ketamine, a comparative analysis was conducted against a control group. These alterations were then validated using the SynGO database in conjunction with the dataset provided by van Oostrum et al. (excel Table S1). (K) Venn diagram analysis, conducted using the FunRich program, was used to visualize the data. (L) Subsequently, enrichment analysis via Enrichr-SynGO of the altered proteome identified significant clustering of these proteins across over 100 synaptic structures. Relevant data are provided in excel Table S2. SynGO biological terms analysis of proteins with decreased (373) and increased (153) abundance. (M and N). (O) Venn diagram analysis, conducted using the FunRich program, was used to visualize the data from Excel Table S4 related to intrinsically disordered proteins (IDPs) found in the decreased and increased proteins of the synaptic structures. All proteasome activity tests are repeated with at least two independent experiments, and each experiment contains four replicates. *Asterisks denote significance levels: *p < 0.05, **p < 0.01, ***p < 0.001. P values were calculated using a two-tailed unpaired t-test to compare the control group with the individual chemical effect. Western blot analyses were performed on independent biological samples with the entire procedure replicated at least twice to confirm consistency of the results.

Initially, we hypothesized that the decrease in p21 levels might be linked to inactivation of the PI3K/Akt pathway, since ketamine and memantine both prompt a rapid and significant reduction in Akt1 phosphorylation (Fig. S1I). Intriguingly, subsequent use of MK-2206, an Akt1 phosphorylation inhibitor at serine 473, and wortmannin, a specific PI3K inhibitor (an upstream kinase of Akt1), revealed that while these agents greatly reduced phosphorylated Akt1 levels, they did not diminish p21 levels as substantially as the NMDAR antagonists (Fig. S1J).

We then extended our investigation to Alzheimer’s disease-IDPs such as Tau and amyloid-beta. We observed that NMDAR antagonists diminished the phosphorylated form of Tau in SH-SY5Y, T98G, and HepG2 cells, and also decreased the levels of constitutively expressed APP695 (beta-amyloid precursor protein) in both Hep3B and SH-SY5Y (Fig. 1C, D).

### NMDAR antagonists enhance proteasome enzymatic activities

The proteasome system is a key protein degradation mechanism in cells ^1-7^. We initially assessed the impact of NMDAR antagonists on proteasome activity by applying fluorogenic substrates to cell lysates using conventional methods ^17, 18^. However, these *in vitro* assays may not accurately reflect the cellular dynamics present in vivo. In line with this, our preliminary experiments indicated that, in cell lysates of HepG2, Hep3B, T98G, and SH-SY5Y, ketamine did not alter the chymotrypsin-, trypsin-, or caspase-like activities of the proteasome (Fig. S2A).

This led us to refine our methodological approach to assess proteasome activity within the context of the cellular environment. We devised a novel ‘In-Cell Fluorogenic Proteasome Assay,’ utilizing cell-permeable substrates derived from an optimized digitonin-tween ratio. Using the in-Cell assay, we found that ketamine significantly enhances the trypsin-like, chymotrypsin-like, and caspase-like activities of the proteasome in a dose-dependent manner across all tested cell lines, including HepG2, Hep3B, T98G, and SH-SY5Y, a result not observed with traditional in vitro methods (Fig. 1E; Fig. S2B, C). We further validated the findings from the traditional cell lysate and in-cell assays using the Cell-Based Proteasome-Glo™ Luciferase System from Promega (Fig. 1F; Fig. S2D).

Subsequent comparative analyses of NMDAR antagonists, ketamine, MK801, and memantine, alongside IU1 ^19^—a known proteasome activity enhancer—and the proteasome inhibitor MG132, revealed that all drugs, with the exception of proteasome inhibitors MG132, upregulated proteasomal activities across various cells to different extents (Fig. 1G; Fig. S2E). Further investigation indicated that proteasome inhibitors MG132 blocked the degradation of p21, p53, and phosphorylated Tau induced by NMDAR antagonists, but not the degradation of phosphorylated Akt1 (Fig. 1H-J, Fig. S2F). Intriguingly, while MG132 predictably inhibited chymotrypsin activity, it unexpectedly increased trypsin activity (Fig. S2G). These observations suggest that NMDAR antagonists may predominantly exert their effects on protein degradation by modulating the proteasome’s chymotrypsin-like activity.

### NMDAR antagonist Ketamine induces changes in synaptic proteins with intrinsically disordered regions in mouse brain tissue

In clinical settings, it has been surprisingly demonstrated that a single intravenous infusion of the NMDAR antagonist ketamine can rapidly alleviate major depression, with effects lasting up to two weeks. Despite extensive research, the full mechanism of action for these drugs remains elusive. Given that disruptions in synaptic function are implicated in neurodegenerative and mental disorders, we sought to investigate ketamine’s effect on synaptic proteins. Building on previous findings that ketamine increases proteasome activities and induces protein degradation, we administered ketamine to mice and harvested brain tissues two hours post-injection for synapse purification and subsequent protein analysis. Of the 1349 synapse-associated proteins quantified, 373 showed a decreased abundance, while 153 showed an increase. In total, 526 proteins exhibited notable changes, as validated against data from SynGO ^20^ and the synaptic protein compendium from van Oostrum et al. ^21^ (Fig. 1K, excel Table S1).

Enrichment analysis via Enrichr-SynGO ^22^ of the altered proteome identified significant clustering of these proteins across over 100 synaptic structures (excel Table S2; Fig. 1L). Importantly, both the decreased and increased protein groups were located similarly, mainly in the synapse, presynapse, postsynapse, membrane (Fig. S2H and I) and enriched similarly with the SynGO-Biological terms (Excel table S3, Fig 1M and N).

Our analysis using the UP_SEQ_FEATURE annotation tool (https://david.ncifcrf.gov/home.jsp) yielded findings that 64.7% of the decreased proteins contained IDRs with a p-value of 5.60E-04 (Fig. 1O 373 excel Table S4)^24^. The term ‘IDR-containing proteins’ did not emerge as significant among the 189 graph records, corresponding to the 823 non-responsive proteins to ketamine, as identified by the SynGO and van Oostrum et al. databases; these proteins did not exhibit significant changes in abundance following treatment (Excel Table S1, S4). A separate analysis conducted with the same UP_SEQ_FEATURE annotation tool on increased proteins yielded findings that markedly contrast with standard expectations. Specifically, 78,1% of these proteins were found to have IDRs, positioning them at the forefront of the enrichment list with a highly significant p-value of 8.30E-08 (Excel Table S4).

Subsequent analysis of protein interactions via the String database (http://www.string-db.org/) ^23^ demonstrated high biological connectivity and non-random associations within each group—increased, decreased, and combined—evidenced by enrichment p-values of <1.0e-16 (Fig. S2J, K).

### Ion channel-specific binding sites modulate NMDAR antagonist effects on proteasome activity

Ionotropic glutamate receptors, including NMDA, AMPA, and kainate receptors, are key to neural transmission^25^. NMDARs differentiate from other ionotropic receptors with their high selectivity for Ca2+ and distinct ligand binding sites for glutamate, glycine, polyamines, channel blockers (e.g., ketamine, memantine, MK801), and allosteric modulators (Fig. S3A).

Examining how different NMDAR sites affect proteasome and p21 protein levels, we found that drugs (Table S1) binding to the channel region, glycine, or specific allosteric sites increased proteasomal chymotrypsin-like activity and reduced p21 levels (Table 1). This effect was absent in polyamines, AMPA and kainate receptors (Table 1, Fig. S3B, C). A summary of the effects on p21 levels and proteasome activity is provided in Table 1, with additional figure representations and detailed analyses omitted for brevity.

**Table 1:** The table displays antagonist and agonist chemicals associated with Glutamate receptors, their binding sites on the receptors, and their effects on proteasome activity and p21 protein levels.

| Name and used dose | Receptor- binding site-<br>function | chym | p21 level |
| --- | --- | --- | --- |
| (+)-MK 801 maleate (20μM) | NMDA - ion channel-antagonist | I | D |
| Memantine hydrochloride (10μM) | NMDA- ion channel- antagonist | I | D |
| Ketamine hydrochloride (100μM) | NMDA - ion channel-antagonist | I | D |
| (+/-)-1-(1,2-Diphenylethyl)piperidine maleate (50μM) | NMDA- ion channel- antagonist | I | D |
| TCS 46b (50μM) | NMDA- NR1A/NR2B<br>selective- antagonist | I | D |
| Eliprodil (30μM) | NMDA-NR2B selective-<br>antagonist | I | D |
| D(-)-2-Amino-4-phosphonobutanoic acid (D-AP4) (50μM) | NMDA- excitatory amino acid-<br>antagonist | I | D |
| N20C hydrochloride (50μM) | NMDA -channel- antagonist | I | D |
| O-Phospho-L-serine (50μM) | NMDA -----antagonist | I | D |
| Felbamate (20μM) | NMDA- glycine- antagonist | I | D |
| Gavestinel (30μM) | NMDA- glycine- antagonist | I | D |
| L-701,324 (5μM) | NMDA- glycine- antagonist | I | D |
| Arcaine sulfate (20-200μM) | NMDA- polyamine- antagonist | NC | NC |
| Ifenprodil hemitartrate (100-800μM) | NMDA- polyamine- antagonist | NC | NC |
| -(R)-3,4-DCPG | AMPA---- antagonist | NC | NC |
| gammaDGG (2-10mM) | AMPA---- antagonist | NC | NC |
| Kynurenic acid (2-40μM) | AMPA/Kainate---- antagonist | NC | NC |
| UBP 301 (1-20μM) | Kainate----antagonist | NC | NC |
| ACPT-II (1-10μM) | Metabotropic----antagonist | NC | NC |
| N-Methyl-D-Aspartic acid (NMDA) (0,2-20mM) | NMDA- ionotropic- <u>agonist</u> | ND | I |
| D-Aspartic acid (2-20nM) | NMDA-endogenous- <u>agonist</u> | NC | NC |
| DL-Glutamic acid monohydrate (20nM) | NMDA-Glutamatergic---<br><u>agonist</u> | I | D |

I, D, NC, ND, and I-D represent an increase in chymotrypsin activity or p21 level, decrease in chymotrypsin activity or p21, no detectable change, not determined, and dose-dependent changes, respectively.

### NMDAR antagonists appear not to impact autophagosome-lysosome-mediated protein degradation

Autophagy, a process previously suggested to be involved in the neuroprotective effect of memantine**^26^**, involves the formation of autophagosomes and autophagic flux, which includes the degradation of microtubule-associated protein light chain 3-II (LC3-II) following fusion with lysosomes **^27^**. Upon autophagy induction, LC3-I is conjugated to phosphatidylethanolamine to form lipidated LC3 (LC3-II). Therefore, the amount of LC3-II correlates well with the number of autophagosomes, and memantine was reported to elevate LC3-II levels **^27^**. In our study, HepG2 cells stably expressing LC3 manifested increased LC3-II post-ketamine treatment (Fig. 2A), hinting at autophagosome accumulation potentially due to autophagy induction. Yet, the total LC3 levels remained unchanged, indicating that autophagic flux, particularly LC3-II degradation upon lysosomal fusion, might not be occurring.

**Fig. 2.**
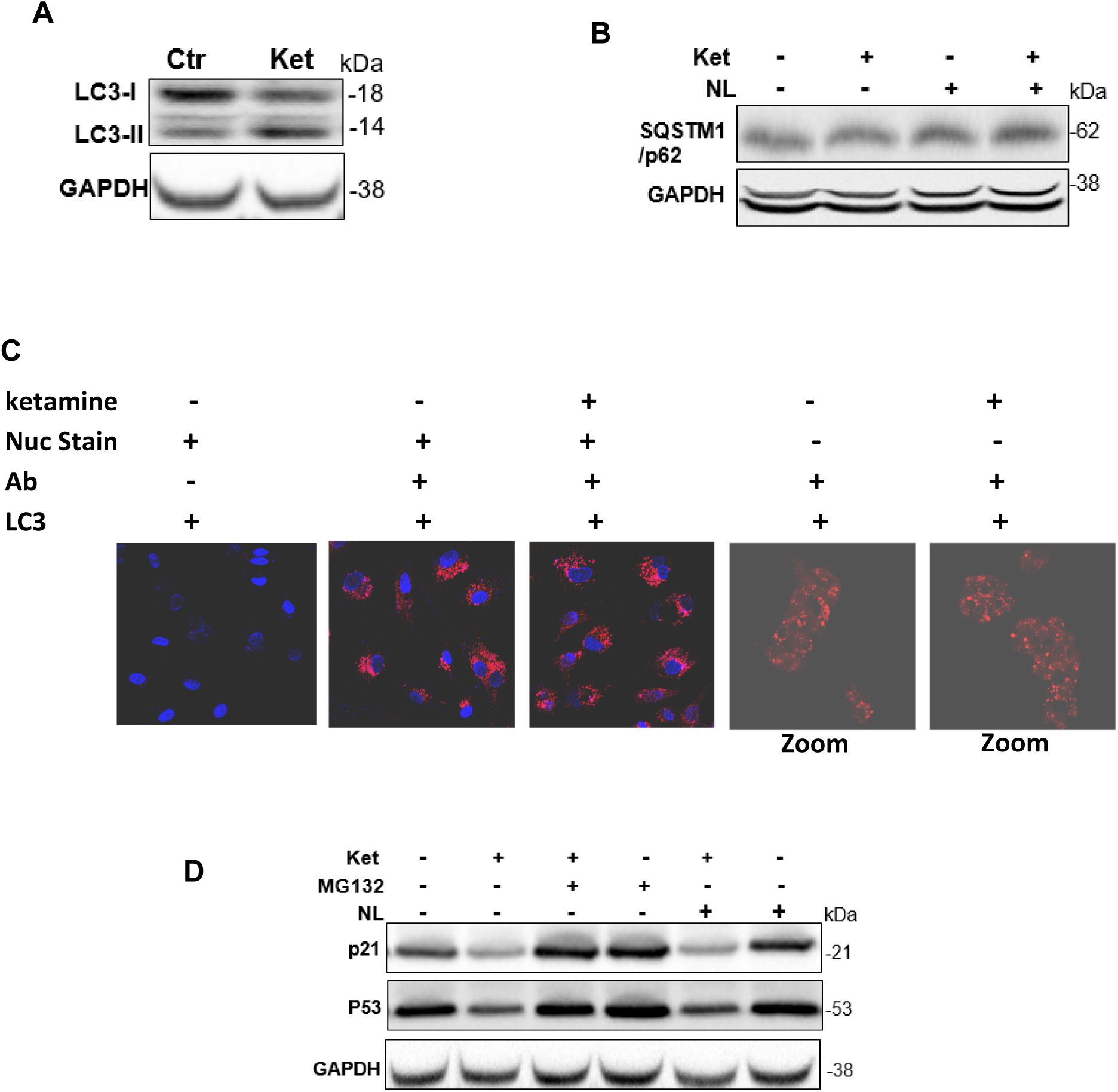
NMDAR blockers’ effects on autophagy-lysosome-related protein degradation pathways. (A) Ketamine’s modulation of LC3-I to LC3-II conversion. (B) p62 protein level alteration in response to ketamine. (C) Confocal visualization of LC3 puncta in response to ketamine versus control in LC3-stable HepG2 cells. (D) Role of NMDA receptor inhibition on p21 and p53 protein degradation with MG132 (proteasome inhibitor) or NL (NH4Cl (20mM) and leupeptin (100µM)) (lysosome inhibitors).

Furthermore, ketamine didn’t affect p62 (SQSTM1) levels nor LC3 puncta density, both markers used to monitor autophagic flux, as evaluated by fluorescent LC3 labeling or immunohistochemistry in LC3-expressing cells ^27^ (Fig. 2B, C). Direct evidence that NMDAR antagonists do not act via the autophagosome-lysosome pathway to degrade p21 and p53 is provided by the fact that proteasome inhibitors, not lysosome inhibitors (NH4Cl and leupeptin), blocked the degradation of these proteins induced by the NMDAR antagonist ketamine (Fig. 2D).

### The ubiquitination pathway is not linked to NMDAR antagonist-mediated proteasome activity

We explored if NMDAR antagonists affect the ubiquitin-proteasome system (UPS), which sequentially employs E1, E2, and E3 enzymes for protein degradation ^3, 5^. To discern if NMDAR antagonists exert their effects via ubiquitin-mediated proteasomal degradation, we developed a model in the Hep3B cell line, utilizing Ub-YFP, a fluorescent reporter that serves as an indicator of 26S UPS activity ^28^. Notably, Ub-YFP is unaffected by 20S UIPS degradation. We identified molecules that are involved in different steps of the ubiquitination process and degradation by the 26S proteasome (Table S2) and targeted them to inhibit Ub-YFP degradation.

Experiments revealed that the E1 enzyme inhibitor PYR-41 inhibited Ub-GFP degradation, as indicated by fluorescence microscopy analysis of YFP expression and proteasome inhibitor, MG132 (Table S2, Fig. 3A). Western blot analysis further confirmed that while PYR-41 led to an elevation in YFP levels within cells, ketamine did not influence YFP levels (Fig. 3b). It is noteworthy that, unlike p21 and p53, YFP— a mutant form of GFP—does not contain any IDRs (Fig. S4).

**Fig. 3.**
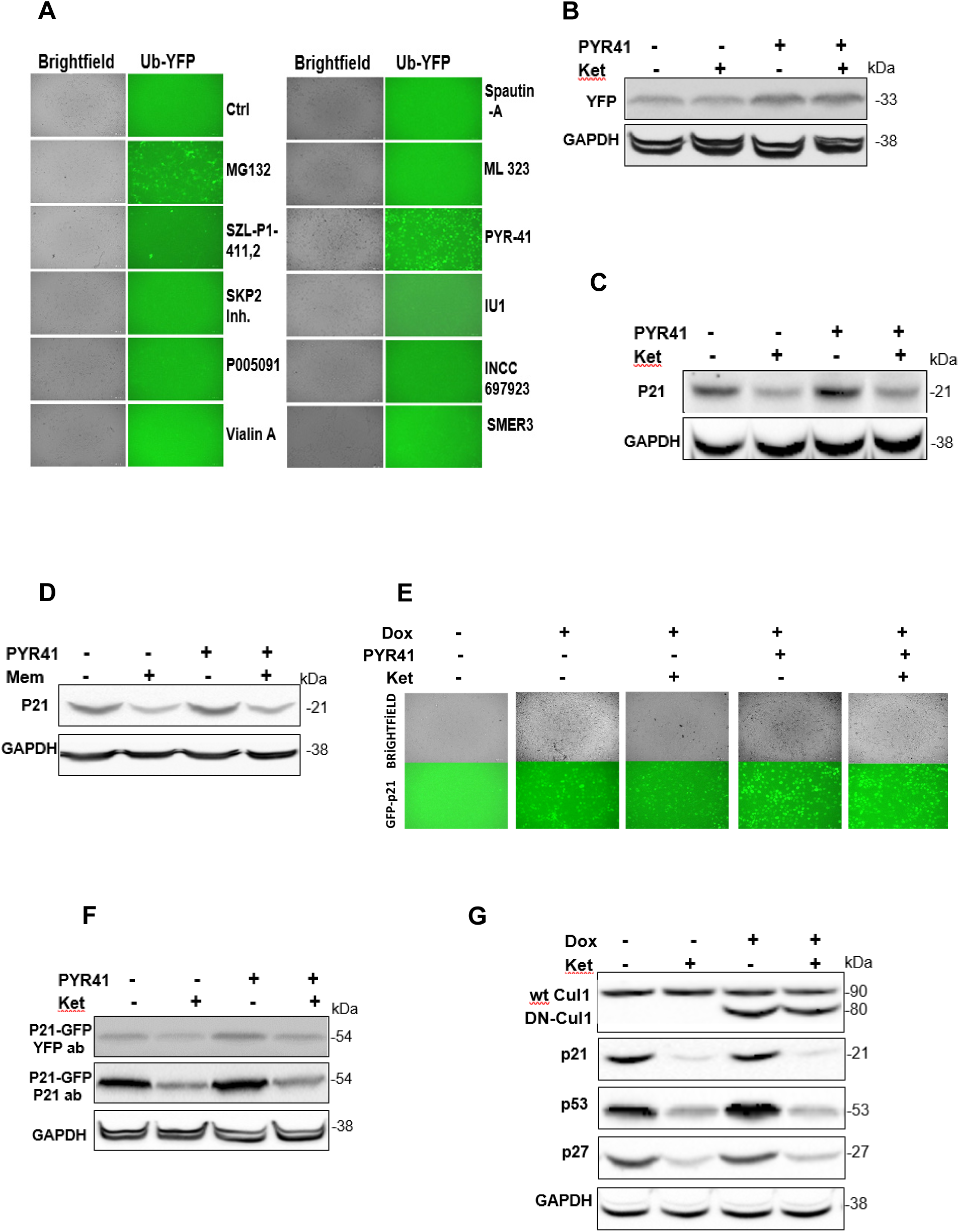
NMDA Receptor Antagonists Do Not Influence Ubiquitination within the Ubiquitin-Proteasome System (UPS) (A) Efficacy of various ubiquitination-related molecules (refer to Supplementary Table S2) on Ub-YFP degradation in Hep3B cells. (B) Impact of PYR-41 (10 µM-12 h incubation), ketamine (500 µM), and their combination on Ub-YFP degradation. (C and D) Modulation of endogenous p21 degradation by PYR-41, ketamine, and memantine in HepG2 cells, both separately and in combination. (E) Differential effects of PYR-41 and ketamine on GFP levels in Hep3B cells with p21-GFP, observed via fluorescence microscopy with Dox induction. (F) Alterations in GFP and p21 levels by ketamine in the presence of PYR-41 in p21-GFP-expressing Hep3B cells, shown by Western blot. (G) Influence of DM-CUL1 on p53 versus p21 and p27 highlighting SCF E3 ligase’s role in ubiquitination, and how ketamine affects p53 degradation in this context.

It is known that the p21 protein is degraded by both the UPS and UIPS ^29^. Investigating further, HepG2 cells—unlike Hep3B, which lack detectable levels of p21—were employed to test the impact of NMDAR antagonists on endogenous p21 in the presence of PYR-41. While PYR-41 increased the p21 protein level by inhibiting its ubiquitination and subsequent 26S proteasome degradation, NMDAR antagonists, ketamine and memantine, reduced p21 levels regardless of PYR-41 treatment, pointing to a ubiquitination-independent pathway (Fig. 3C, D). Similarly, when p21 was tagged with GFP in Hep3B cells, NMDAR antagonists still lessened p21-GFP levels despite the inhibition of the ubiquitination process (Fig. 3E, F).

E3 ubiquitin ligases confer specificity to the UPS by selecting substrates for ubiquitination. With hundreds of E3 ligases identified in humans, the SCF complex—comprising Skp1, Cul1, Rbx1, and an F-box protein—is a notable E3 type. Cul1 serves as a scaffold for the complex, facilitating substrate recognition and ubiquitination **^30, 31^**. To probe SCF’s involvement in protein ubiquitination, we utilized a dominant-negative Cullin 1 (DM-CUL1) construct to disrupt SCF activity in HepG2 and T98G cells. Despite the successful expression of DM-CUL1, levels of p21 and p27 remained unchanged, signaling these proteins are not SCF substrates. However, DM-CUL1 expression markedly increased p53 levels, an SCF ligase substrate, but this increase was annulled by ketamine treatment, demonstrating that NMDAR antagonists can still degrade p53 despite SCF ligase inactivation in both cell lines (Fig. 3G). The reduction of p53 was more pronounced in the presence of NMDAR antagonists compared to when SCF ligase activity was unperturbed.

Collectively, these results indicate that the UPS is not primarily involved in the proteasome-mediated degradation of p21 and p53 proteins induced by NMDAR antagonists. The experiments suggest that such antagonists may engage alternative proteolytic mechanisms, as shown by the effective degradation of proteins irrespective of ubiquitination status.

### Regulators of the 19S subunit of the 26S proteasome do not mediate the effects of NMDAR antagonists on proteasomal activity

Ubiquitinated proteins are targeted by the 19S regulatory subunit of the 26S proteasome, which encompasses non-ATPase and ATPase components such as Rpn1, Rpn2, Rpn10, Rpn13, among others. To assess NMDAR antagonist interactions with these 19S subunit regulators, we first examined how these antagonists affect proteasome-mediated degradation and chymotrypsin-like activity in the context of Rpn13 inhibition. Rpn13, along with Rpn10 and Rpn1, is crucial for recognizing ubiquitinated substrates within the 19S subunit ^5^. We used RA190, a selective Rpn13 inhibitor, and noted that p21 levels rose upon RA190 application, likely attributed to impeded UPS-mediated degradation (Fig. 4A). Furthermore, ketamine effectively degraded p21 whether or not cells were pre-treated with RA190. We also found that NMDAR antagonists consistently enhanced chymotrypsin-like activity of the proteasome with or without RA190 intervention, as visualized in a line graph for hourly measurements and a bar graph for the 15th-minute mark (Fig. 4B and C). Concurrently, the abundance of ubiquitinated proteins increased in the presence of RA190, yet ketamine application led to a decrease in their levels (Fig. 4D).

**Fig. 4.**
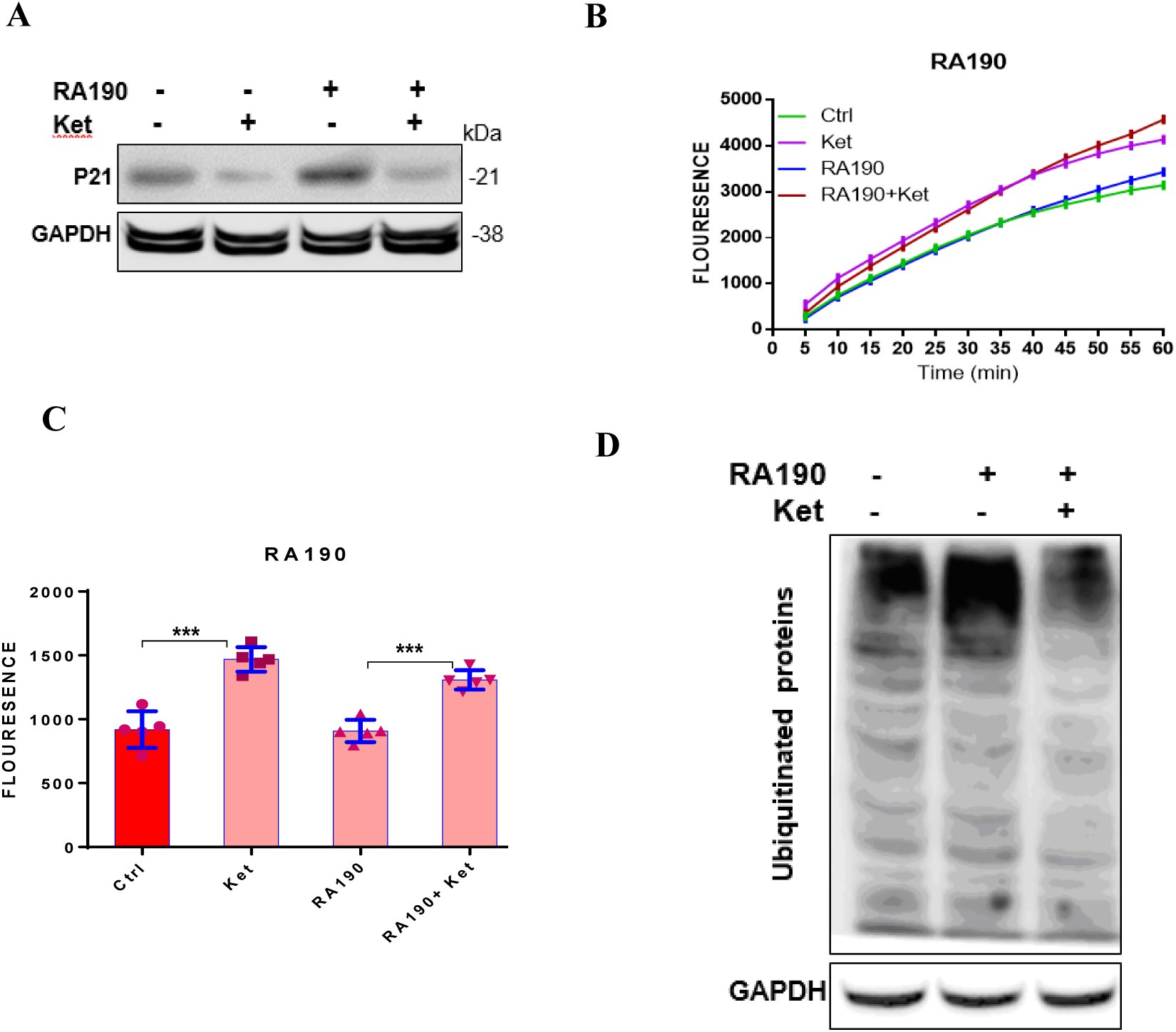
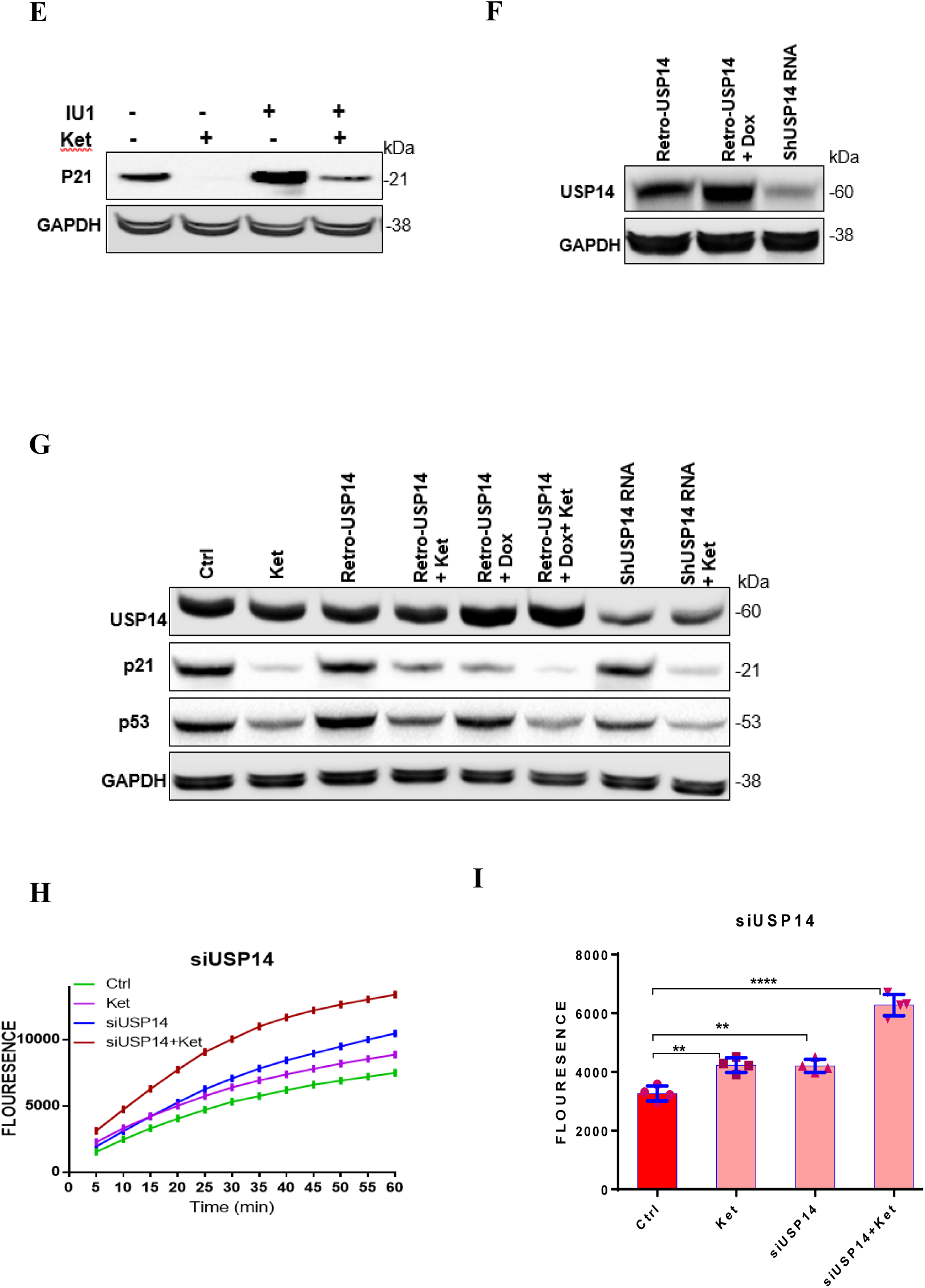
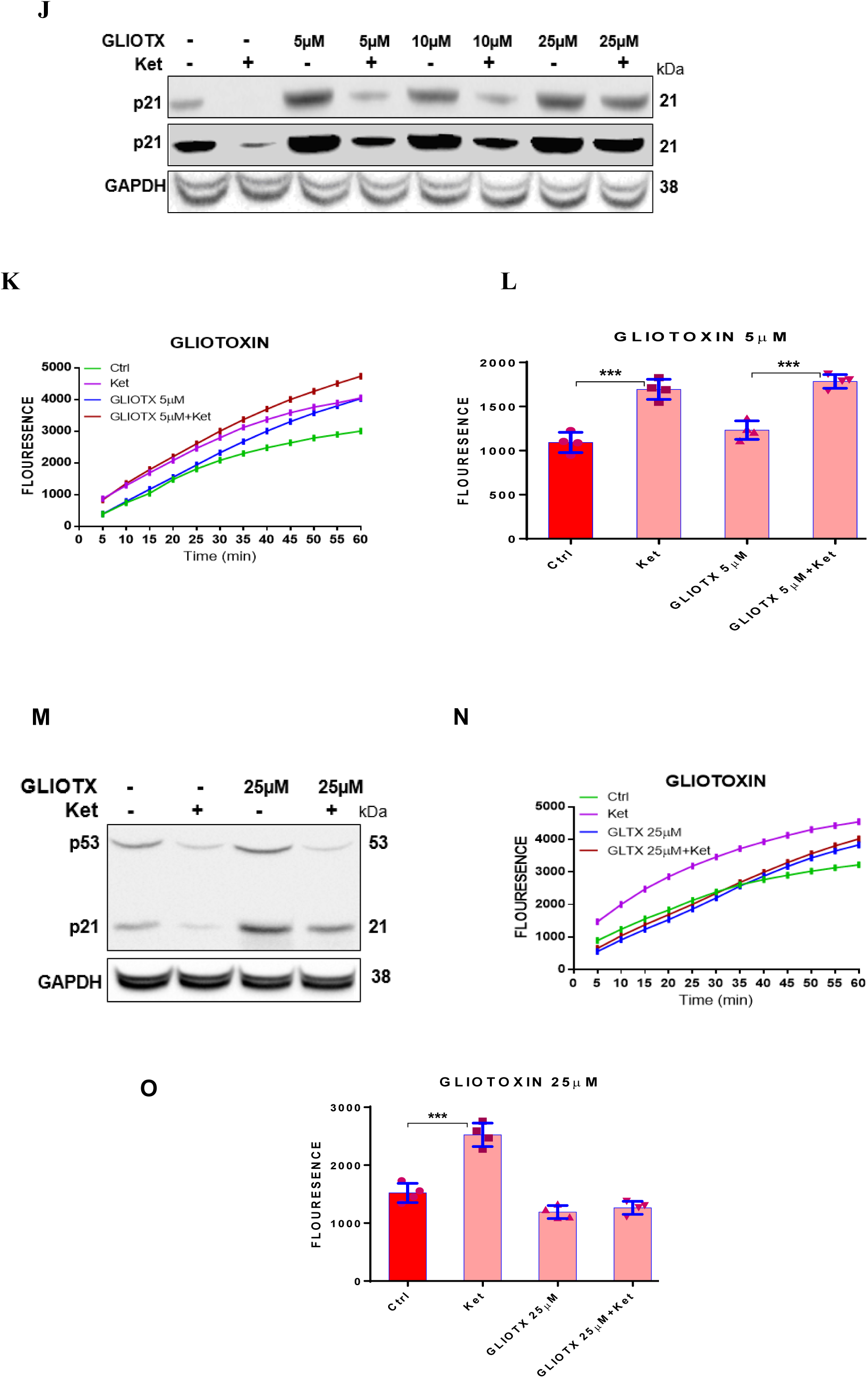
NMDAR antagonists do not impact the regulatory particles of the 19S subunit of the 26S proteasome. (A) Effects of RPN13 inhibition by RA190 on p21 levels and the corresponding influence of ketamine in RA190 (1µM, 12 h incubation) pre-treated conditions. (B and C) Role of RA190 in chymotrypsin-like activity and ketamine’s intervention in these conditions. (D) RA190’s elevation of ubiquitinated proteins contrasted with ketamine’s reduction effect. (E) IU1’s effect on p21 protein levels and the modifying role of ketamine alongside USP14 inhibition. (F) Changes in USP14 protein levels with overexpression or silencing techniques. (G) How USP14 expression levels affect p21 and p53 proteins and ketamine’s influence under these conditions. (H, I) Impact of USP14 expression alterations on chymotrypsin-like activity and modifications by ketamine. (J) Dose-dependent effects of the PSMD14 inhibitor gliotoxin on p21 levels were examined, alongside the effects of ketamine on p21 with and without gliotoxin pre-treatment, with evidence that gliotoxin inhibits 20S proteasome chymotrypsin activity at high doses. p21 levels were analyzed via western blot at two time points of signal exposure. (K and L) Ketamine’s effects on chymotrypsin activity were evaluated with and without 5 µM gliotoxin. (M) At 25 µM gliotoxin, ketamine fails to decrease p21 levels while reducing p53 levels. Due to this discrepancy, the Western blot was repeated multiple times and the result was ultimately confirmed by using p21 and p53 antibodies concurrently on the same membrane (N and O) Chymotrypsin activity in response to ketamine was assessed with and without 25 µM gliotoxin. Results are presented as the mean ± SEM, based on 3 to 5 technical replicates from each of n = 3 independent experiment. *Asterisks denote significance levels: *p < 0.05, **p < 0.01, ***p < 0.001. P values were calculated using a two-tailed unpaired t-test to compare the control group with the individual chemical effect or USP14 knockdown cells.

The aforementioned experiments with ketamine and memantine were conducted on at least two different cell lines, specifically HepG2 and T98G, to assess their effects on the levels of p21 and p53 proteins, as well as on chymotrypsin-like activity. However, only results pertaining to the impact of ketamine on p21 levels and chymotrypsin activity are presented in the following sections, except in cases where notable differences emerged.

The 19S regulatory particle encompasses various deubiquitinases (DUBs) that perform deubiquitination within the ubiquitin-proteasome system (UPS). Key DUBs associated with the 19S unit include the USP family’s USP14, the JAMM family’s POH1 (also known as RPN11/pda1/S13/mpr1), and the UCH family’s UCHL5 (UCH37)^5^. USP14 is a critical enzyme; IU1, a specific inhibitor of USP14, is one of the few known compounds that can enhance the proteasome’s chymotrypsin-like activity in vivo^19^. Our findings confirm that IU1 significantly raises both chymotrypsin-like and caspase-like activities of the proteasome (Fig. 1J). Notwithstanding, unlike NMDAR antagonists, IU1 does not lower but rather substantially increases p21 levels. In contrast, NMDAR antagonists consistently decrease p21 levels, even in the presence of IU1 (Fig. 4E).

To verify and elucidate the impact of NMDAR antagonists on DUB function, particularly regarding USP14, we employed a lentivirus-based silencing approach with USP14-specific siRNA and, conversely, upregulated USP14 expression using a doxycycline-inducible retroviral system (Fig. 4F). Knockdown of USP14, in contrast to the effects of IU1, did not alter p21 levels (Fig. 4G). However, silencing USP14 led to a decrease in p53 levels (Fig. 4G). Interestingly, ectopic expression of USP14 decreased p21 abundance. Notably, the reduction of p21 and p53 levels in USP14-silenced cells in the presence of ketamine was comparable to, if not greater than, the reduction observed in control cells treated with ketamine. Importantly, USP14 silencing increased chymotrypsin-like activity similarly to the USP14 inhibitor IU1. Furthermore, NMDAR antagonists augmented chymotrypsin activity more dramatically in USP14-silenced cells than in controls (Fig. 4H and I). These findings, in line with those concerning the ubiquitin ligase SCF, propose that the degradation of p21 and p53 via the UPS involves distinct molecules. Therefore, although the mechanisms underlying p21 and p53 degradation by the UPS vary, the capacity of NMDAR antagonists to diminish the levels of these proteins appears to be independent of the UPS-related processes investigated thus far.

In subsequent experiments, we tested the effects of b-AP15 and N-ethylmaleimide, both of which inhibit UCHL5 and USP14, similar to IU1. These compounds exhibited comparable impacts to IU1, as illustrated in the figures S5A, B, and C (the results for b-AP15 are not shown).

In a subsequent study, we investigated gliotoxin, recognized as an inhibitor of the DUB family member RPN11, also known as PSMD14 ^32^. Gliotoxin is also known to inhibit the chymotrypsin-like activity of the 20S proteasome at higher concentrations ^33^. Treatment with gliotoxin led to increased p21 levels, suggestive of inhibited UPS-mediated degradation (Fig. 4J). Notably, NMDAR antagonists degraded p21 efficiently at gliotoxin doses of 5 and 10 µM, comparable to the untreated control group (Fig. 4J). Moreover, NMDAR antagonists elevated chymotrypsin-like activity of the proteasome to the same or greater extent in the presence of gliotoxin than without it (Fig. 4K, L). At a higher gliotoxin dose of 25 µM, p21 degradation was reduced in the presence of NMDAR antagonist (Fig. 4J, M). Intriguingly, p53 levels decreased similarly in cells treated with the high dose of gliotoxin and in the control group (Fig. 4M). Further, a 25 µM dose of gliotoxin also inhibited chymotrypsin-like activity of the proteasome in the presence of NMDAR antagonists (Fig. 4N, O). These findings suggest that while NMDAR antagonists impact p21 degradation by modulating chymotrypsin-like activity, different mechanisms, potentially independent of chymotrypsin-like activity such as trypsin or caspase like activities, appear to govern p53 degradation by the proteasome.

Parallel experiments were conducted using capzimin, another PSMD14 inhibitor, yielding results akin to those observed with low doses of gliotoxin (Fig. S6A-C).

Additionally, the outcomes of PSMD14 knockdown were consistent with the effects of both gliotoxin and capzimin, thereby corroborating the previous findings and offering more targeted insights (Fig. S7A-D).

The collective findings from the described experiments suggest that the action of NMDAR antagonists is independent of the ubiquitination and deubiquitination processes within the UPS.

It is noteworthy that the inhibition of DUBs increased the chymotrypsin-like activity of the proteasome. However, this did not result in the degradation of ubiquitinated proteins, such as p21 and p53; rather, there was an accumulation within the cell, since the removal of their ubiquitin chains by DUBs is a prerequisite for their processing through the UPS. Importantly, a synergistic increase in chymotrypsin-like activity was observed when cells were treated simultaneously with NMDAR antagonists and DUB inhibitors, compared to the effect of each agent alone. This suggests separate mechanisms of chymotrypsin activity induction within the 26S and 20S proteasome components. The assay for chymotrypsin-like activity, utilizing the Suc-LLVY peptide substrate, does not require deubiquitination, thus allowing for the detection of increased activity within the 26S proteasome system, even in the presence of DUB inhibitors such as IU1 and the other inhibitors used in this study. Therefore, it appears that DUB inhibitors activate chymotrypsin-like activity at the 19S regulator of the 26S proteasome, while NMDAR antagonists specifically enhance the activity within the 20S core.

### NMDAR Antagonists Activate the 20S Proteasome

Oxidatively modified proteins, known to be highly susceptible to degradation, have been shown to be preferentially targeted by the 20S proteasome ^34, 35^. In addition, whereas the 26S proteasome’s activity is diminished in the presence of H_2_O_2_ at concentrations as low as 400 μM and completely inactivated at 1 mM, the 20S proteasome maintains its function, unaffected by H_2_O_2_ concentrations up to 5 mM ^34,35^.

In light of these results, we devised a set of experiments to not only validate the previously shown findings but also to consolidate our experimental data. We established the concentrations at which H_2_O_2_ impedes degradation mediated by both UPS (26S proteasome) and UIPS (20S proteasome), using cells expressing p21-GFP, and we assessed the effects of NMDAR antagonists (Fig. S8A). H_2_O_2_ caused a slight increase in GFP levels at 400 µM, with a more substantial increase observed at 4 mM (Fig. S8A). We observed a striking increase in endogenous p21 levels in HepG2 cells exposed to 800 µM H2O2, leading us to hypothesize that this concentration primarily inhibits 26S proteasome activity, thereby causing an accumulation of ubiquitinated p21 protein due to the impairment of 26S proteasome-mediated degradation (Fig. 5A; Fig. S8B). When we investigated the effects of ketamine on p21 levels in cells exposed to 800 µM H_2_O_2_, we observed that ketamine significantly reduced p21 levels to a greater extent in these cells compared to its effect on p21 levels in control cells (Fig. 5A; Fig. S8C). This trend was paralleled in the overall levels of ubiquitinated proteins (Fig. 5B). These findings suggest that NMDAR blockers may not only facilitate the degradation of ubiquitinated proteins but also appear to be more effective in degrading ubiquitinated proteins than non-ubiquitinated proteins. We recorded even greater increases in p21 levels with 4 mM H2O2, which, we postulated, completely inactivated the 26S proteasome and significantly impeded the 20S proteasome activity (Fig. 5C). Notably, at this higher concentration of H_2_O_2_, NMDAR antagonists lost their capacity to degrade p21 (Fig. 5C). A similar pattern was observed for the levels of ubiquitinated proteins examined in the context of 4 mM H_2_O_2_ treatment (Fig. 5D).

**Fig. 5.**
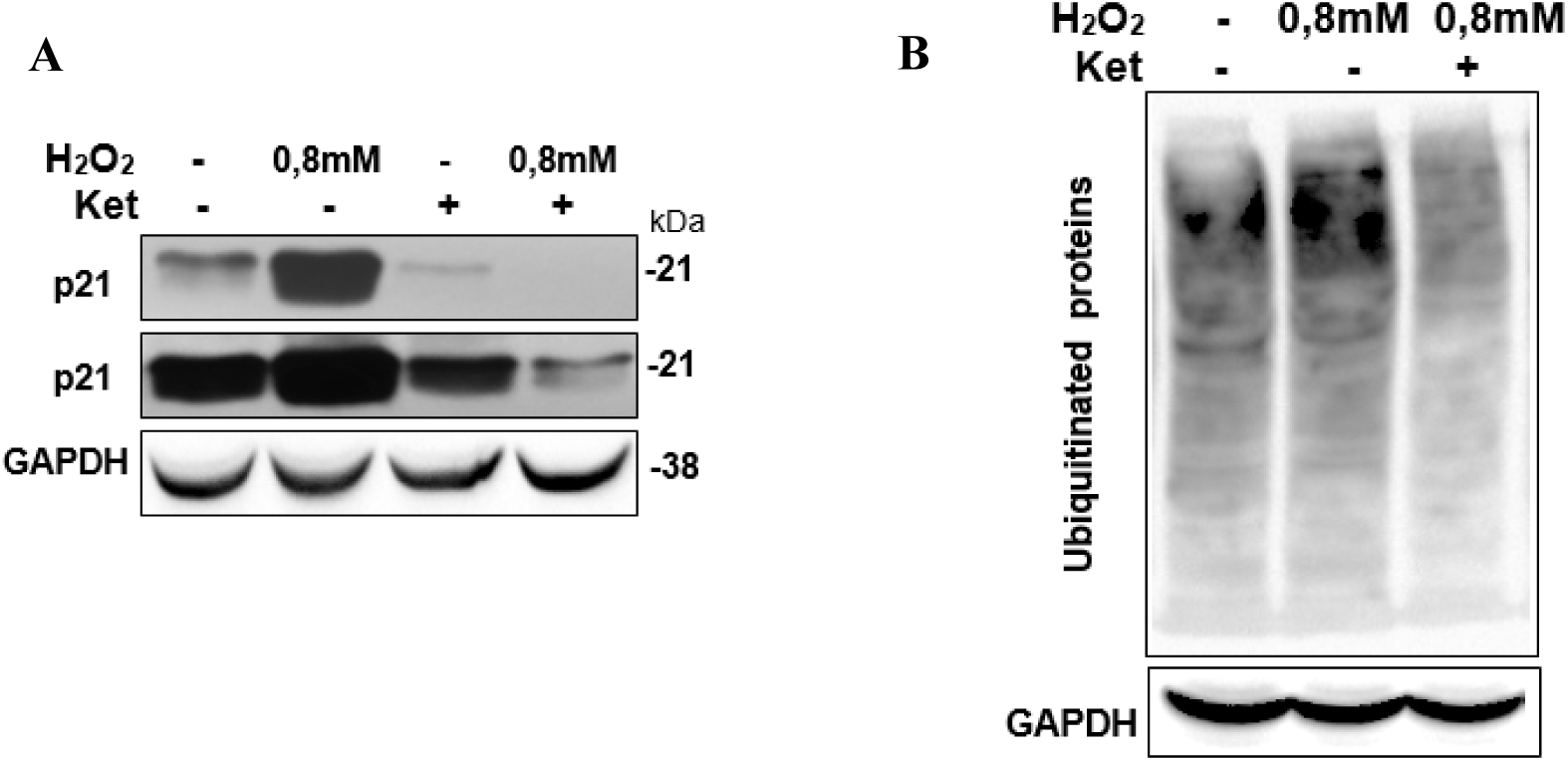
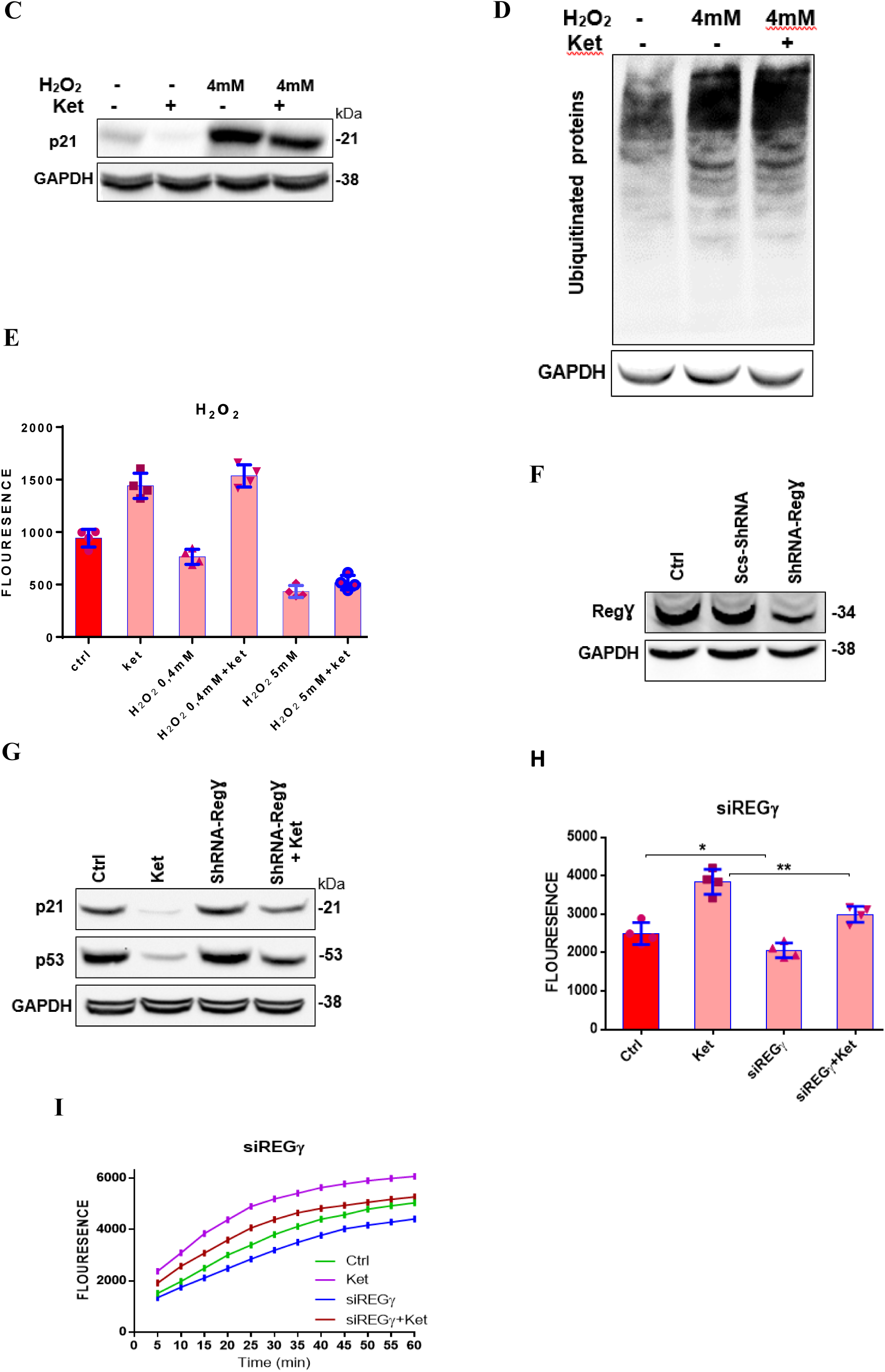
NMDAR Antagonists modulate 20S Proteasome activities. (A) Oxidative impact of 0.8mM H_2_O_2_ on p21 protein levels and ketamine’s effect under these circumstances. (B) Effects of 0.8mM H_2_O_2_ on ubiquitinated proteins and changes due to ketamine exposure. (C) 4mM H_2_O_2_’s influence on p21 levels and ketamine’s subsequent effects in such oxidative conditions. (D) How 4mM H_2_O_2_ alters ubiquitinated protein levels and the corresponding ketamine response. (E) Examination of H_2_O_2_ at various concentrations on chymotrypsin activity and ketamine’s role in such scenarios. (F) Effects of silencing REGγ in HepG2 cells. (G) Results of ketamine on p21 and p53 levels in REGγ-silenced cells versus controls. (H, I) Impact of ketamine on chymotrypsin activity in REGγ-silenced and control cells. Results are presented as the mean ± SEM, based on 3 to 4 technical replicates from each of n = 3 independent experiment. *Asterisks denote significance levels: *p < 0.05, **p < 0.01, ***p < 0.001. P values were calculated using a two-tailed unpaired t-test to compare the control group with the individual chemical effect or REGγ knockdown cells.

The observed changes in chymotrypsin-like activity, tested in the presence of H_2_O_2_, ketamine, or both, are consistent with and elucidate the protein level results. Specifically, proteasomal chymotrypsin-like activity was reduced in the presence of both 0.4 mM and 5 mM H_2_O_2_. While ketamine enhanced proteasomal chymotrypsin-like activity at 0.4 mM H_2_O_2_, this enhancement was not observed at 5 mM H_2_O_2_ (Fig. 5E).

Subsequent experiments were aimed at demonstrating that NMDAR antagonist activity is directly linked to the function of the 20S proteasome. Non-ATPase particles, such as the 11S complex (consisting of PA28 α, β, and γ), are known to associate reversibly with the 20S proteasome and regulate its activity. PA28γ (REGgamma), in particular, is recognized for its role in binding and activating the 20S proteasome ^36^. In the following experiments, PA28γ expression in HepG2 and T98G cells was knocked down using lentivirus-delivered siRNA specific to PA28γ (Fig. 5F). Cells with silenced PA28γ exhibited reduced p21 and p53 degradation in response to NMDAR antagonists as compared to the control cells (Fig. 5G). Further, the overall proteasome activity, as well as its increase in response to NMDAR antagonists, was diminished in the PA28γ-silenced cells relative to control cells (Fig. 5H, I). These results underscore that the activity of NMDAR antagonists is primarily exerted on the 20S proteasome, and specifically implicate PA28γ, a regulatory component of the 20S system, in the NMDAR antagonist-induced enhancement of proteasomal function.

We also demonstrate that the NMDAR antagonist, ketamine, does not directly activate the purified 20S proteasome (devoid of regulatory proteins), nor does it enable the purified 20S proteasome to degrade His-tagged p21 protein (expressed in and purified from Hep3B cells) (Fig. S9 A, B) Moreover, ketamine treatment did not induce significant changes in the structure of the 26S or 20S proteasome complexes, as determined by Western blotting using an anti-α1−α7 antibody cocktail or native gel electrophoresis (Fig. S9C, D). Furthermore, no substantial alterations were observed in the abundance of several 20S and 26S regulatory proteins co-immunoprecipitated with the α1−α7 subunits in the presence of ketamine (Fig. S9E). These results may suggest that 20S regulatory particles are essential for the 20S proteasome’s functional activity

### Ketamine induces changes in synaptic proteins associated with neurodegeneration

Enrichment analysis, incorporating 526 synaptic proteins categorized by biological processes, pathways, cellular components, and sequence features, was conducted using the DAVID bioinformatics resource. This analysis revealed over 1,000 records significantly enriched with terms pertinent to neurodegenerative conditions and mental health disorders, all of which exhibited strong P values (excel Table S5).

Further refinement of the 526 proteins using enrichment analysis within the pathway category, conducted via the ZS Revelen-PPI tool ^37^ (https://zs-revelen.com), identified terms encompassing a variety of biological mechanisms. These included signal transduction both within and outside synaptic contexts, nervous system development, the neuronal system, membrane trafficking, and neurotransmission. Specific pathways such as tyrosine kinases (TrkB), MAPK family signaling cascades, WNT signaling, and BDNF (Brain-derived neurotrophic factor) were also highlighted (Fig. 6A). Complementary analysis using Enrichr-KG, which covers the Mammalian Phenotype (MP), GO Biological Processes, and KEGG pathway Browser categories, indicated significant enrichment in areas such as hyperactivity, abnormal CNS synaptic transmission, altered excitatory post-synaptic currents, attenuated long-term potentiation, nonsense-mediated mRNA decay, cytoplasmic translation, protein targeting to membrane, ribosomal function, oxytocin signaling pathway, and endocytosis (Fig. 6B). Crucially, the terms identified across the various categories and through the different analytical platforms have all been associated with the pathogenesis of neurodegenerative diseases in prior research ^38-44^.

**Fig. 6.**
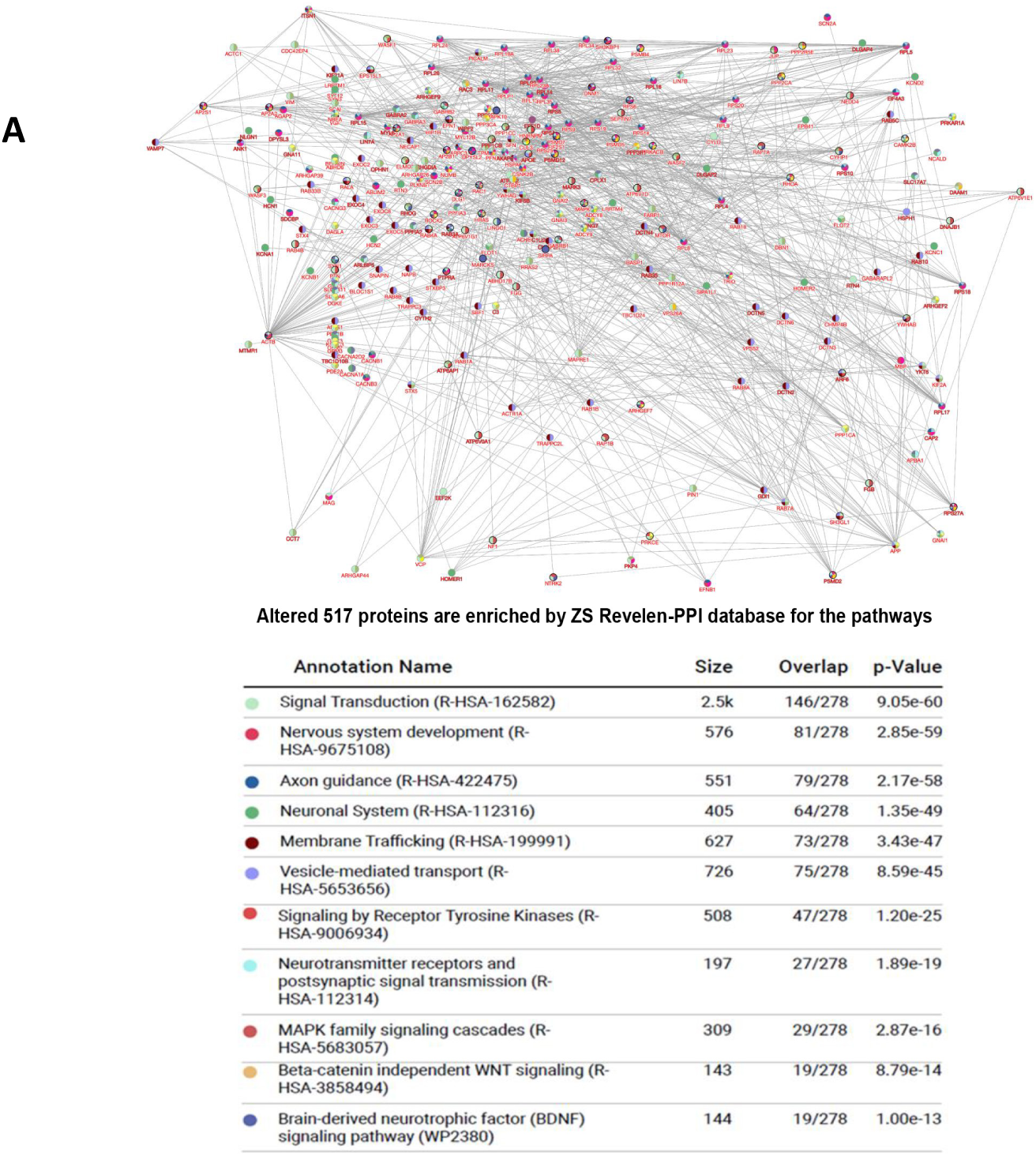
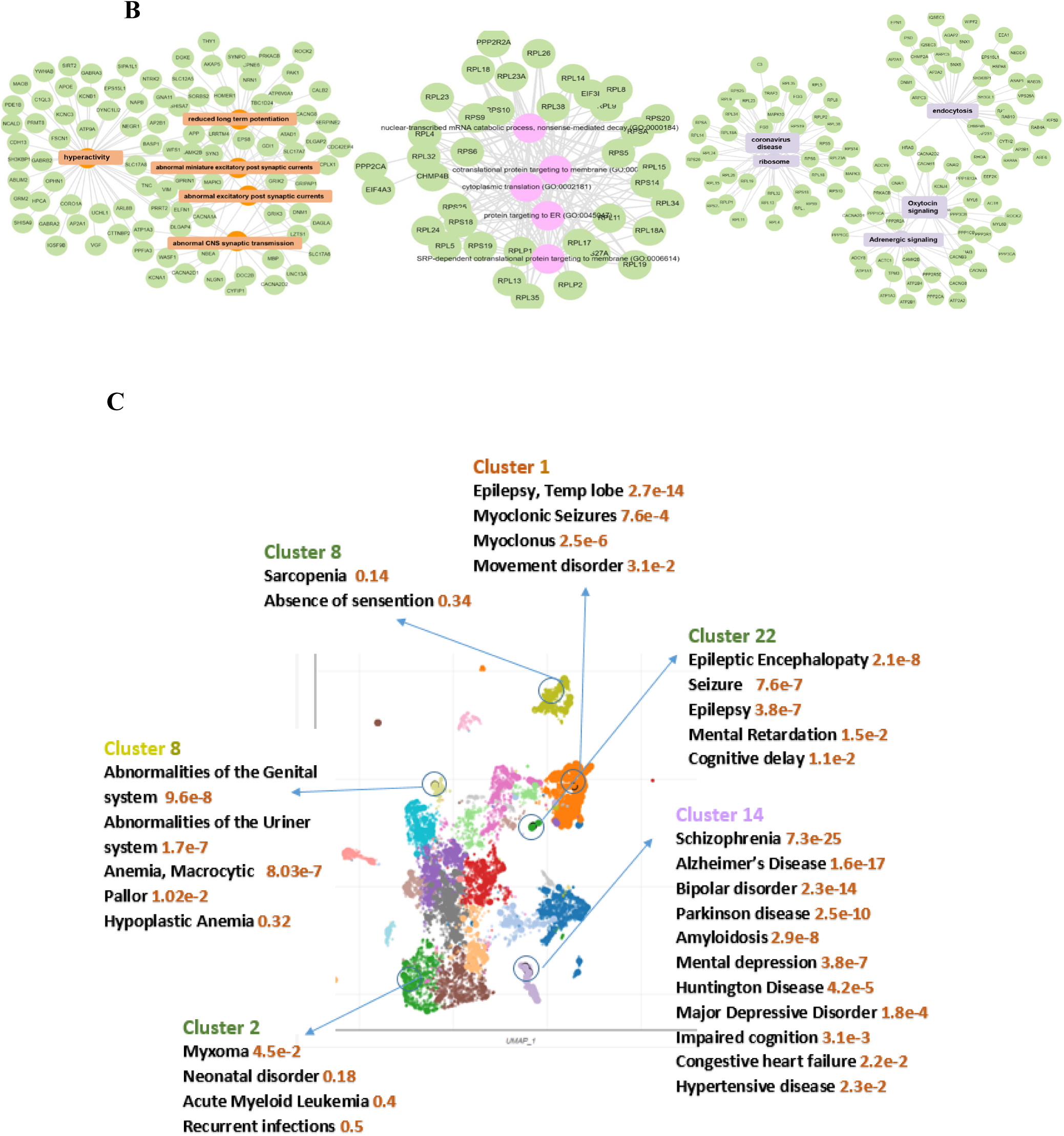
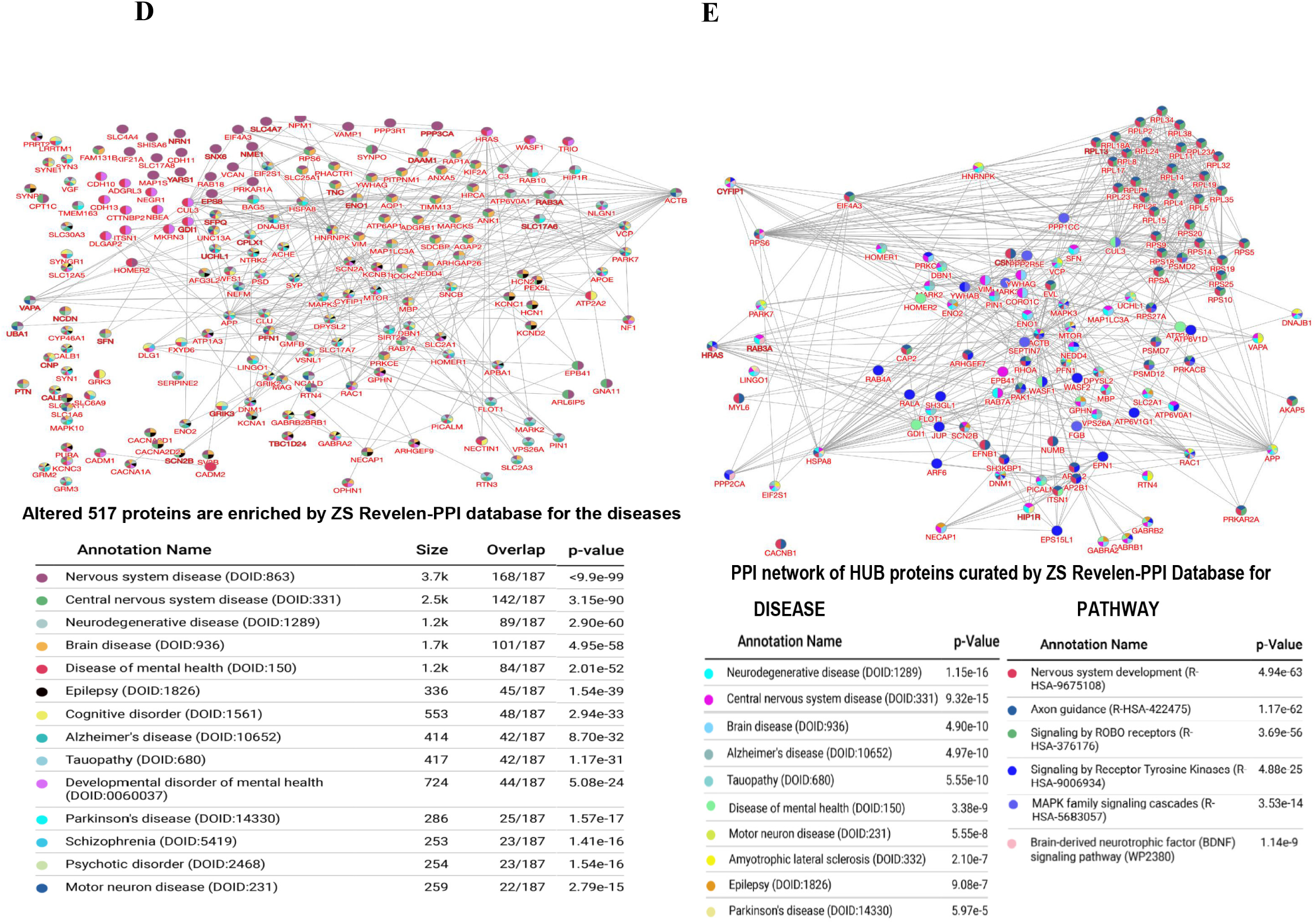
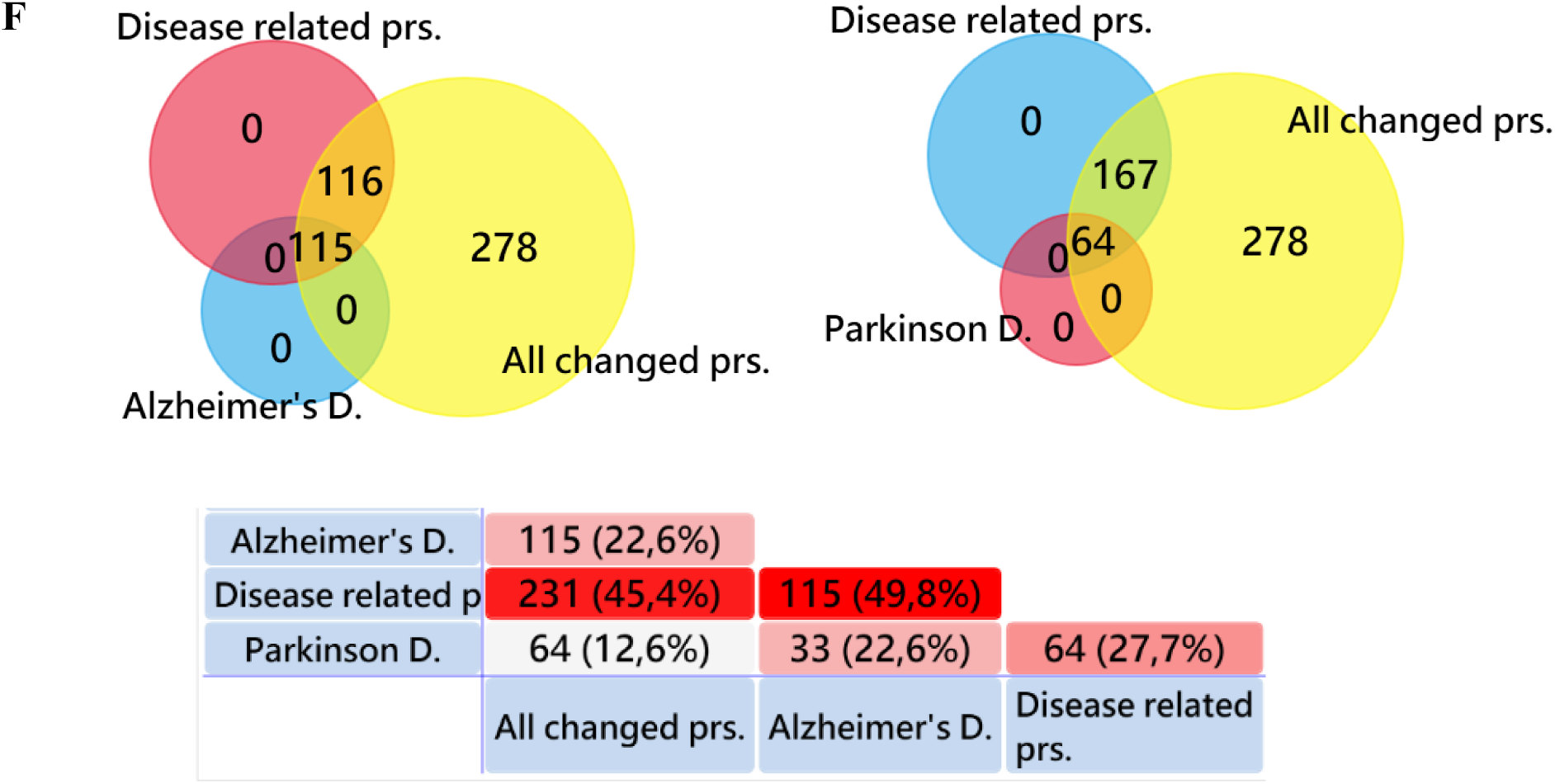
Enrichment of neurodegeneration-related proteins altered by ketamine. (A) Enrichment analysis of 526 proteins in the pathway category via ZS Revelen-PPI indicates top terms and PPI data. (B) Ranking of enriched categories from Enrichr-KG analysis, including Mammalian Phenotype, GO Biological Processes, and KEGG pathway browser. (C) Scatterplot of 526 proteins against DisGeNET disease clusters reveals 25 groups, with significance of top six clusters emphasized. (D) ZS Revelen-PPI tool’s evaluation of the 526 proteins within disease categories. (E) Identification of HUB proteins from the 526-protein set using ZS Revelen-PPI, categorized by disease. (F) Comparative analysis of proteins linked to AD, PD, and 231 disease-associated proteins within the 526 altered set via FunRich Venn (excel Table S8). Some proteins listed in the data were not recognized by the FunRich program.

A scatterplot visualization generated by DisGeNET-Enrichr, a disease association library, was used to examine the 526 proteins. This analysis demarcated 25 distinct disease clusters. Notably, Cluster 14 had the most significant P value among all clusters, encompassing mental disorders such as Schizophrenia and Bipolar Disorder, as well as neurodegenerative conditions like Alzheimer’s Disease (AD), Parkinson’s Disease (PD), and Huntington’s Disease (HD) (Fig. 6C). In a similar vein, the protein-protein interaction (PPI) network for the 526 proteins, construed through the ZS Revelen-PPI tool for disease correlation, underscored a preponderance of nervous system diseases, neurodegenerative disorders, and mental health issues (Fig. 6D).

Within this protein network, proteins that were shown to interact with five or more other proteins were classified as HUB proteins (excel Table S7). An examination of 178 such HUB proteins from the cohort of 526 modified proteins using the ZS Revelen-PPI tool brought to light enrichments closely associated with nervous system diseases, neurodegenerative diseases, brain diseases, and mental health disorders. This reflects the enrichment patterns observed with the entire dataset of 526 proteins (Fig. 6A, B, D and E).

Further analysis using the DisGeNET database within the Enrichr platform ^22^ indicated that 231 out of the 526 proteins are implicated in diseases such as Schizophrenia, Alzheimer’s Disease, Bipolar Disorder, Epilepsy, and Parkinson’s Disease (PD). Notably, Venn diagram analysis conducted by the FunRich program (http://www.funrich.org) ^45^ revealed that approximately 49.8% of the proteins associated with these diseases (231 proteins) are implicated in Alzheimer’s Disease (AD) (excel Table S8, Fig. 6F, and Fig. S10A).

The affected proteins’ enrichment in the KEGG pathway for AD demonstrates connections to most AD etiologies, including mitochondrial dysfunction, proteasome system aberrations, amyloid beta formation, tau hyperphosphorylation, autophagy dysregulation, impaired axonal transport, and beyond (Fig. S10B). Similarly, for PD, the altered proteins are involved in pathways known to be disrupted by PD pathology (Fig. S10C).

### Ketamine induces changes in synaptic proteins associated with mental diseases

It has been determined that major depressive disorder symptoms improve within two hours after a single intravenous infusion of the NMDAR antagonist ketamine, with effects lasting for up to two weeks. However, the mechanism by which ketamine alleviates these symptoms and sustains its effects for such an extended period remains unknown. Recent research increasingly recognizes that dysfunctional synaptic changes, including synaptic transmission, plasticity, and stability, play a crucial role in the pathology of depression and other mood disorders ^14, 46-49^. In agreement with this perspective, enrichment analyses of the proteins modulated by ketamine, conducted using DAVID bioinformatics Functional Annotation and SynGO 2022—which assesses ontology, localization, and biological processes—significantly highlight synaptic-related terms (excel Table S2, S3, and S6).

In a pivotal analysis, a Volcano plot generated from the 526 proteins using DisGeNET within the Enrichr framework brought schizophrenia and Bipolar Disorder to the forefront as the diseases most connected to the protein alterations (Fig 7A). This was further substantiated by Venn diagram analyses performed with the FunRich tool, which showed that 65.4% of the disease-associated proteins (231) have linkages with schizophrenia and Bipolar Disorder (excel Table S8, Fig 7B, and Fig. S11A)

**Fig. 7.**
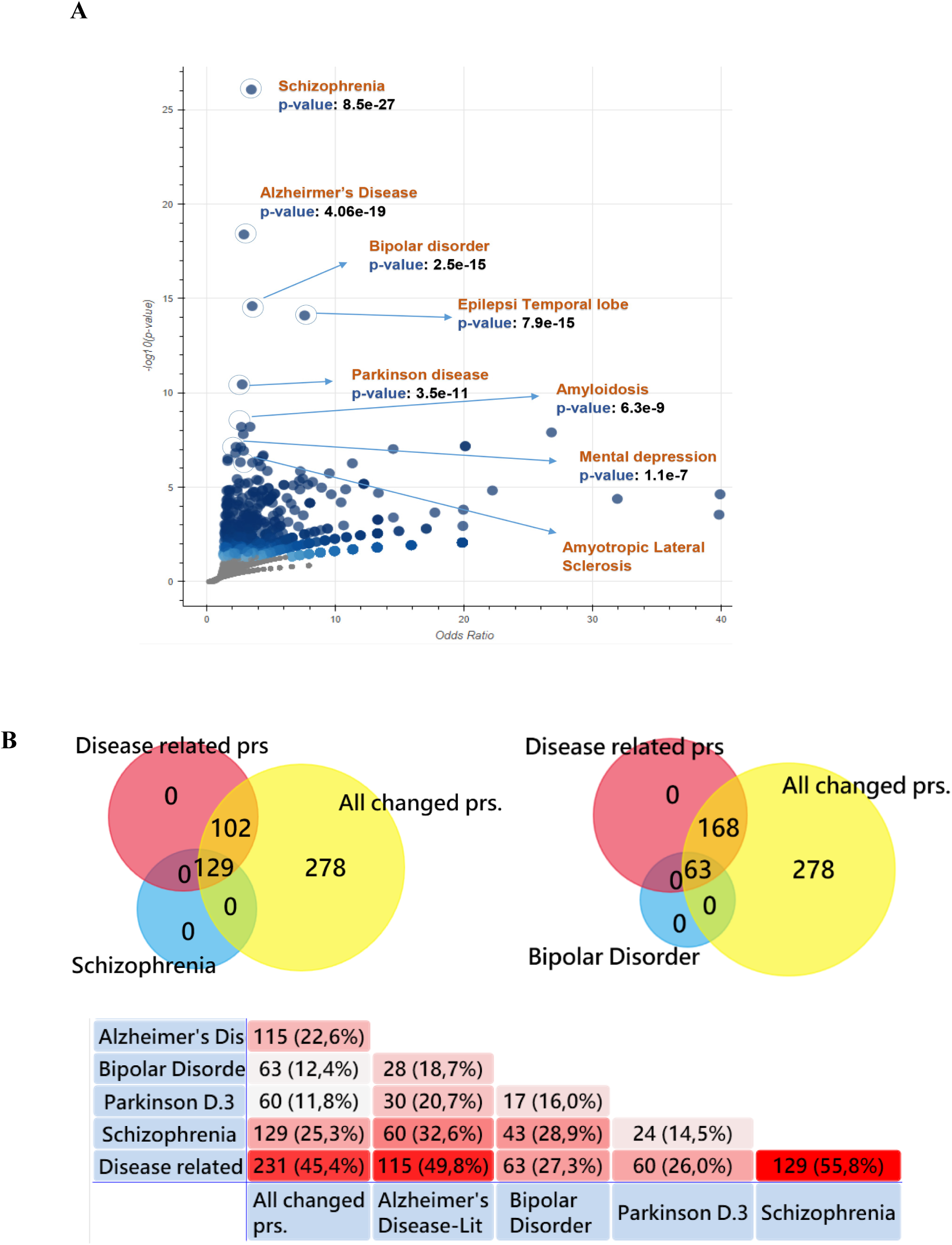
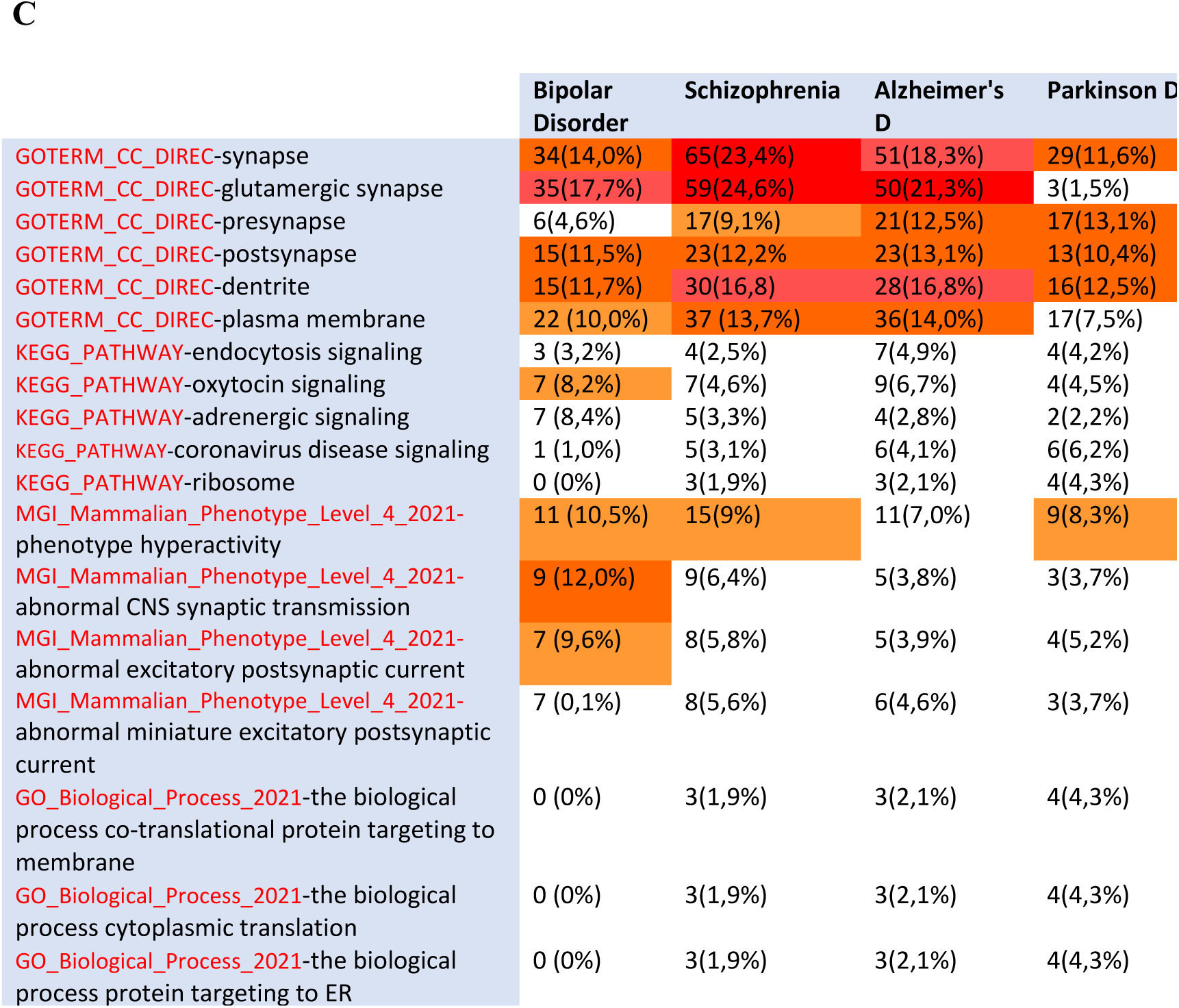
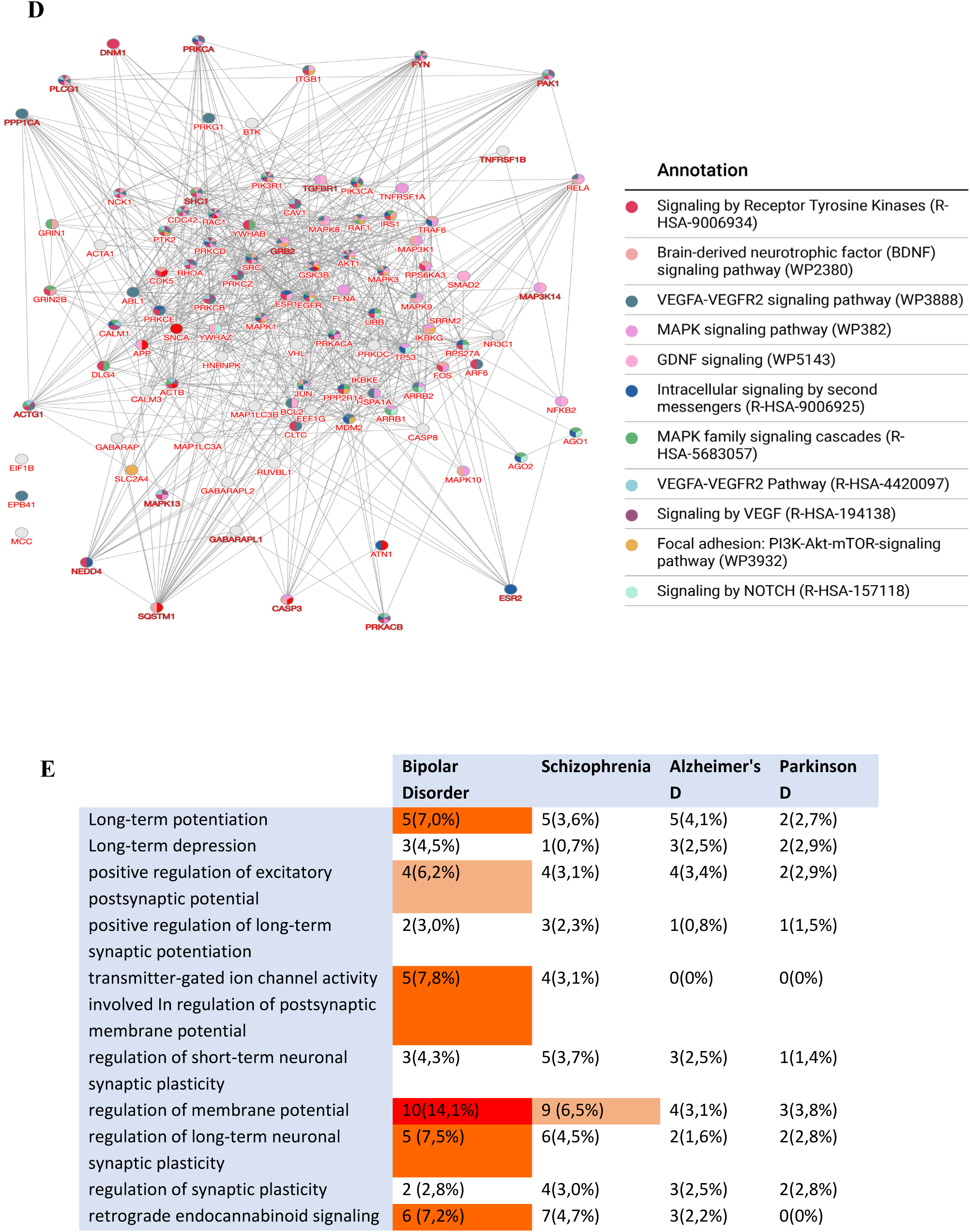
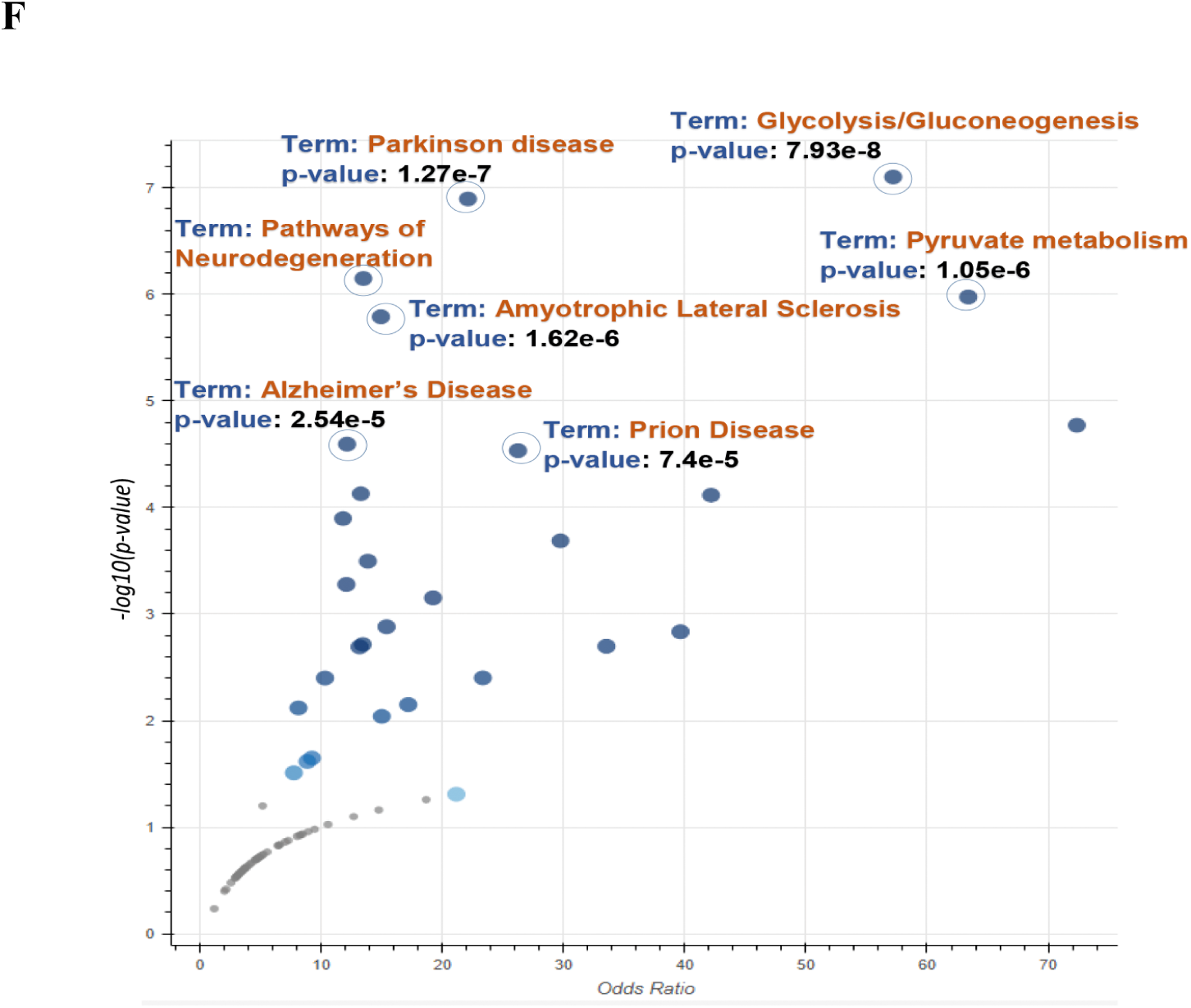
Ketamine induced protein alterations in synaptic function linked to mental disorders. (A) Volcano plot analysis via DisGeNET shows schizophrenia and bipolar disorder most associated with the 526 altered proteins.(B) Venn diagram analysis investigates overlaps between schizophrenia and BD within 231 disease-associated proteins from the 526 altered set. Pairwise comparisons for AD, PD, schizophrenia, and BD are detailed in Dataset S7. (C) FunRich Venn compares proteins linked to schizophrenia, BD, AD, and PD across biological processes, pathways, and cellular locations. (D) PPI network analysis for HUB proteins from a refined 372-protein subset via the ZS Revelen-PPI tool for Pathways category. (E) FunRich Venn examines proteins related to schizophrenia, BD, AD, and PD within synaptic plasticity terms, highlighting proteins most abundant in individual terms.(F) Enrichr-Volcano analysis of EC-related proteins referenced from O’Donovana, S. M. et al.

Alterations in glutamatergic neurotransmission at the synapse have been implicated in the etiology of schizophrenia, depression, and Alzheimer’s disease. Notably, Parkinson’s disease (PD) is often excluded from this association ^50-52^. Within our study, pairwise comparison via FunRich Venn analysis reveals that among the 526 proteins identified as altered, there is a pronounced representation within the glutamatergic synapse in the context of proteins associated with schizophrenia, Alzheimer’s Disease (AD), and Bipolar Disorder (Fig. 7C). Moreover, our results suggest a limited impact of NMDAR antagonists on the glutamatergic synapse in PD compared to the aforementioned conditions. The data shows that while over 20% of proteins at the glutamatergic synapse are affected in schizophrenia, BD, and AD, a mere 1.5% of PD-related proteins exhibit alterations (Fig. 7C).

Previous studies have revealed that ketamine’s antidepressant effects, similar to conventional antidepressants, are closely linked to BDNF-TrkB signaling^53-57^. Consistent with this, our DAVID bioinformatics analysis of all modified proteins showed significant enrichment in the neurotrophin (BDNF) and TrkB, along with MAPK, AMPA-Glutamate, and Akt-mTOR signaling pathways (excel Table S6). Furthermore, analysis of the protein-protein interaction (PPI) network of the 526 proteins, organized by pathways using the ZS Revelen-PPI tool, and the identification of HUB proteins revealed prominent terms associated with nervous system development and signaling pathways involving TrkB, MAPK, and BDNF (Fig 6A and E). Similarly, enrichment of the top 20 HUB proteins drawn from the dataset of 526 proteins, conducted using PPI HUB protein tools from the Enrichr platform, underlined pathways related to neurotrophins, MAPK, long-term potentiation, and neurodegeneration (Fig. S11B; excel Table S9). Most notably, subset analysis of the 375 proteins with decreased expression further emphasized the importance of neurotrophic factors such as BDNF, VEGF, and GDNF, along with TrkB and MAPK signaling pathways (Fig. 7D). Thus, our findings contribute to the growing body of evidence that underscores the significant link between BDNF-TrkB signaling and the molecular mechanisms underlying depression.

### Ketamine causes changes in plasticity and potentiation-related proteins

BDNF’s versatility is highlighted by its role in various adaptive neuronal processes, including long-term potentiation (LTP), long-term depression (LTD), specific kinds of short-term synaptic plasticity, and the homeostatic regulation of intrinsic neuronal excitability ^58-59^. Consequently, researchers have posited that the long-term effect of ketamine on depressive symptoms can be attributed to its action on synaptic plasticity, particularly homeostatic plasticity ^58, 59^. An examination of plasticity and potentiation-related terms within the DAVID Annotation chart for the 526 proteins altered by ketamine revealed over 18 pertinent terms, such as ‘Long-term potentiation’ and others (excel Table S6 and S10). Notably, ‘retrograde endocannabinoid signaling’—recognized as a crucial regulator of both short- and long-term plasticity ^60^—emerged as a significant theme from the DAVID enrichment analysis (Fig. S11C). Moreover, a FunRich Venn pairwise comparison underlined a more substantial association of BD with these terms, in contrast to schizophrenia, AD, and PD (Fig. 7E). These findings align with previous research that links ketamine to synaptic plasticity, contributing to its persistent effects on major depressive symptoms.

Recent studies suggest that ketamine is not rapidly eliminated but remains bound to NMDAR in areas such as the lateral habenula, and this extended inhibition may be responsible for plasticity-related alterations and the sustained impact of NMDAR antagonism ^61^. However, our results show that although the NMDAR antagonist reduces p21 and other proteins within just 10 minutes, the levels of all proteins decreased rebound to baseline between 12 to 24-hours post-treatment (Fig. S1E).

Electroconvulsive Therapy (ECT) remains the most potent acute intervention for severe depression that is often resistant to treatment. So far, meta-analyses exploring its mechanisms have implicated plasticity processes that follow ECT, such as neurotrophic factors (particularly BDNF), neurogenesis, dendritogenesis, and synapse formation ^62^. In research examining the impact of ECT on the hippocampal proteome, significant alterations in metabolism-associated proteins were observed after both acute and chronic ECT treatments ^63^. Upon applying DAVID bioinformatics and SynGO tools to proteins identified in a referenced study ^63^, we found that synapse-related proteins were also altered, despite synaptic protein purification not being part of the experimental protocol (Fig. S11D, E). Crucially, when analyzed for enrichment within the KEGG pathway and DisGeNET databases, terms emerged such as epilepsy, PD, AD, Amyotrophic Lateral Sclerosis (ALS), and neurodegeneration pathways — findings that closely parallel those from ketamine administration (Fig.7F).

## Discussion

While neurodegenerative diseases are characterized by a multitude of intricate etiologies, deficiencies within the proteasome systems emerge as a prominent contributing factor to their onset ^10-12, 40-44^.

Notably, proteasome functionality has been observed to decline with advancing age. This decline correlates with the abnormal accumulation of IDPs, and/or misfolded proteins, which have been implicated in the pathogenesis of neurodegenerative diseases ^6-8^.

Evidence from various studies suggests that the 20S proteasome (UIPS) plays an indispensable role in the clearance of IDPs, IDPRs, and proteins that are oxidatively damaged or misfolded ^1-4, 6-8^. Therefore, there exists significant therapeutic interest in the development of strategies aimed at augmenting the in vivo activity of the 20S proteasome system, with the aim of delaying, preventing, and/or treating aging related disorders.

In this study, we presented that NMDAR antagonists can potentiate proteasome activities by means of ubiquitin/ATP-independent pathways directly through the 20S proteasome. This breakthrough extends our understanding of protein degradation pathways, revealing that the 20S proteasome can degrade ubiquitinated proteins, potentially with a higher efficiency under some conditions. A variety of experiments, encompassing the inhibition of Rpn13, a regulator of the 19S subunit, suppression of DUB regulatory proteins (USP14, PSMD14) of the 19S using multiple drugs or knockdown techniques, and the comprehensive inhibition of 26S proteasome activity by H_2_O_2_, have revealed that NMDAR antagonists may operate independently of the entire 26S proteasome complex, including the 19S regulatory component. It is noteworthy that the 20S proteasome exists independently of the 26S complex and is abundant in cells, comprising approximately 1% of the total cellular protein**^6^**. Notably, agents such as IU1 that enhance chymotrypsin-like activity and inhibit USP14 within the 19S regulatory particle did not impact protein degradation, thereby also underscoring the essential role of the 20S proteasome’s autonomous proteolytic activity.

It is evident from our investigation that not all examined proteins are susceptible to degradation by the 20S proteasome system. This observation underscores the existence of a selective prerequisite governing the degradation of proteins by the 20S proteasome ^64^. The proteomic analysis of synaptic proteins revealed that ketamine markedly alters the synaptic protein profile, predominantly affecting those identified as disordered.

The alterations in synaptic plasticity-related mechanisms, encompassing neurotrophic factors (such as BDNF and others), MAPK, postsynaptic AMPA signaling, and synapse formation, play a pivotal role in the pathogenesis of various mental diseases ^51-57^. In this study, we demonstrate that the impact of NMDAR antagonists on these processes mirrors the effects observed with classical antidepressant MAO inhibitors and ECT.

Our results reveal a rapid onset of protein reduction, initiated as early as 10 minutes following the administration of NMDAR antagonists (Fig. S1E). This reduction persisted for a brief period, lasting 8-12 hours in cell cultures, before returning to baseline levels (Fig. S1F). The swift replacement of damaged or non-functional proteins with functional counterparts within such a short timeframe may elucidate the rapid and pronounced therapeutic effects of ketamine on mood disorders.

Changes in various forms of synaptic plasticity have been suggested as the underlying mechanism for the long-term effects of ketamine in major depression ^58-60^. Our results notably highlight that BD, in comparison to PD, AD, and Schizophrenia, exhibits a significantly higher enrichment in terms associated with plasticity and potentiation. This enrichment encompasses retrograde endocannabinoid signaling, a pivotal pathway governing both short- and long-term plasticity (Fig. 7e).

Despite the primary application of NMDAR antagonists in neurodegenerative diseases such as AD and PD, our results consistently highlight schizophrenia as the leading condition in each of the enrichment analyses. Moreover, our enrichment analysis indicates that ketamine induces changes in sets of proteins enriched in over 1,000 disease chart records. These records cover a diverse range of conditions, including neoplasia, various forms of anemia, diabetes, and numerous other diseases, with over 200 specifically associated with brain diseases (excel Table S5).

Therefore, our findings are important not just for AD and PD, but they also suggest new treatment options for many different diseases linked to proteins that don’t fold properly or are affected by various causes.

## Author contributions

Conceptualization and designing the experiments: FS

Performing the Investigations: FS, AG, BTA, UG, BS, IU, MT, OC, SK

Research expenses: FS

Project administration: FS

Supervision: FS

Writing – original draft: FS

## Data and materials availability

All data available in main text or the supplementary materials.

## MATERIALS AND METHODS

### Cells

We used HepG2 and Hep3B hepatocellular carcinoma, T98G glioblastoma, and SH-SY5Y neuroblastoma cell lines. They were cultured in DMEM with 10% FBS, 1% Pen/Strep, and 1% L-glutamine, and maintained at 37 °C with 5% CO2

### Antibodies

We employed a range of antibodies targeting p21, p27, Akt1, phospho-Akt1, Ubiquitin, USP14 (Cell Signaling Technology); p53, Serine racemase, GFP, APP695, PA28γ, PA28α, PA28β, MECL-1, LMP2, SQSTM1/p62, 20S Proteasome subunits, GAPDH, beta-actin (Santa Cruz Biotechnology); Tau, phospho-S199 Tau (Thermo Fisher Scientific); LC3 (Sigma Aldrich); PSMD14, Rpn2, PSMD4, Rpn13 (Solarbio), and YFP (Biovision), all in line with the providers’ usage guidelines.

### Chemicals and Kits

Various chemicals and kits were utilized: MG132, RA190, Gliotoxin, Capzimin, (+/-)-1-(1,2-Diphenylethyl) piperidine maleate, TCS 46b, Eliprodil, AP4, N20C hydrochloride, O-Phospho-L-serine, L-701,324, D-Aspartic acid, Kynurenic acid, UBP 301, (R)-3,4-DCPG, gammaDGG, DL-Glutamic acid monohydrate, L-Glutamic acid monosodium salt (Santa Cruz); Lactacystine, Ketamin hydrochloride, Suc-LLVY-AMC, P005091, Vialin A, Spautin A, ML 323, SKPIN C1, SZL-P1-411, SMER3, SKPIN C1, NCC697923, PYR41, IU1, Lactacystin (Cayman Chemical); Memantine hydrochloride, dizocilpine maleate (MK-801), Wortmannin, MK2206, cycloheximide (Sigma-Aldrich); Z-LLE-AMC, Ac-RLR-AMC (AdipoGen Life Sciences); b-AP15 (MedChemExpress), and Proteasome luciferase system (Promega); (+)-MK 801, Felbamate, Gavestinel, N-Methyl-D-Aspartic acid, ACPT-II (Tocris Bioscience); Leupeptin, N-Ethylmaleimide, rProteinA/G beads, Ni-NTA Ultra-Flux Affinity Microsphere (Solarbio), and 20S Proteasome Assay Kit (Enzo Life Sciences). Memantine, Ketamine, and MK801 concentrations varied as specified, typically 0.01-0.1mM, 0.1-1mM, and 0.1-0.2mM in vitro. Manufacturers’ recommendations were followed for all items.

### Western Blot Analysis

Cell lysates were prepared in a buffer with 20 mM Tris (pH 7.5), 150 mM NaCl, 1 mM EDTA, 1 mM EGTA, 1% Triton X-100, and protease inhibitors (Cell Signalling Technology). Fifty micrograms of protein were separated on 8%-12% Bis-Tris gels (Invitrogen) and blotted onto nitrocellulose membranes (Bio-Rad). After blocking with 5% skim milk in PBS-Tween for 1.5 hours, membranes were incubated with primary antibodies overnight at 4°C.

### Proteasome activity assay using cell lysate

To assay proteasome activity in cell lysates, 2 million cells were lysed using glass beads at 4°C with a 50 mM Tris–HCl (pH 7.5), 5 mM MgCl2, 1 mM DTT, and 250 mM sucrose buffer. Protein concentrations of the supernatants were determined using BCA protein assay (Pierce). Samples of 10 micrograms of protein were added to a 96-well plate with 26S proteasome assay buffer and fluorogenic substrates (Suc-LLVY-AMC, Z-LLE-AMC, Ac-RLR-AMC) for chymotrypsin-like, caspase-like, and trypsin-like activity, respectively. The resulting AMC fluorescence was tracked with a Biotek fluorometer at 5-minute intervals for 60 minutes at 37°C.

### Proteasome activity assay using “In-Cell Fluorogenic Proteasome Assay”

We developed cell-permeable versions of fluorogenic substrates for a non-lysate living cell assay to measure proteasome activity effectively. In a 96-well plate format, 8×10^3 cells per well were seeded. After 24 hours, the medium was changed to include an NMDAR blocker. Two hours later, 100 µl of 2X fluorescence cell buffer was added (1 ml 1M Tris-HCl pH:7.5, 500 µl 1M KCl, 200 µl 1M NaCl2, 200 µl 0.1M MgCL2, 50 µl 0.2% Tween20, 75 µl digitonin 20mg/ml in DMSO ChemCruz cat no: sc-280675, 10 µl fluorescence substrates including Suc-LLVY-AMC, Z-LLE-AMC, and Ac-RLR-AMC Ac-RLR-AMC, 8.1 ml H2O). After incubation at room temperature for 5 minutes, the fluorescence of each sample was measured using a fluorescence spectrometer (Biotek instrument -Synergy HT).

### Use Promega’s Cell-Based Proteasome Luciferase Systems to assess proteasome activity in living cells

To measure proteasome activity in live cells, Promega’s Proteasome-Glo™ Luciferase System was used. We seeded 8,000 cells per well in a 96-well plate, and after 24 hours, treated them with an NMDAR blocker. Two hours later, 100 μl of the prepared Proteasome-Glo™ Reagent was applied per well. Post a 10-minute room temperature incubation, the luminescence was recorded using a Biotek luminometer

### Preparation of P21-GFP cDNA and Cloning into Doxycycline-Inducible Retroviral Vector

The pBSK-clo2D vector, previously constructed in our lab to include multiple cloning sites (KpnI, ApaI, XhoI, SalI, PmeI, BamHI, EcoRV, EcoRI, XbaI, NotI, HindIII, MfeI, EcoRI, PstI, SmaI, BamHI, XbaI, BstXI, SacI), was utilized to create a GFP-p21 fusion. The GFP open reading frame was PCR-amplified from pEF-GFP (courtesy of Connie Cepko, Addgene plasmid #11154) using primers GFP-F and GFP-R (sequences: ATCCACCGGTCGCCACCATG and CTTGTACAGCTCGTCCATGC). The resulting fragment was cloned into the EcoRV site of pBSK-clo2D, which had been linearized and blunt-ended using Phusion High-Fidelity DNA Polymerase (Thermo Scientific).

After linearization with EcoRV, the pBSK-clo2D-GFP vector was dephosphorylated using Alkaline Phosphatase (Fermentas). Separately, RNA was extracted from HepG2 cells using the RNase MiniKit (Qiagen) and reverse-transcribed using random primers (Invitrogen) and M-MLV reverse transcriptase (Invitrogen) to generate cDNA. The p21 gene was amplified via PCR with specific primers (sense, 5’-CAT GTC AGA AAC CGGCTG GGG-3’; anti-sense, 5’-TTA GGG CTT CCT CTT GGA GA-3’) and subsequently cloned into the dephosphorylated vector using T4 DNA Ligase (Thermo Scientific)

The fusion of GFP and p21 was designed to be in-frame. To create the P21-GFP fusion construct, the related fragment was excised from pBSK-clo2D using BamHI, and then ligated into the BamHI site of a pre-treated pRetroX-Tight-Pur vector. Successful constructs were amplified in DH5α E. coli.

For the inducible expression system, HepG2, Hep3B, and T98G cell lines were first integrated with the rtTA transgene using rtTA-containing viral particles produced in Hek293T cells. These cells were transfected with pRetroX-Tet-On-Advanced and the helper vectors pCMV-VSV-G, pUMVC, using the CalPhos Mammalian Transfection Kit and culturing under polybrene. Post-infection, cells underwent selection with neomycin (G418) for 10 to 15 days, after which survivors were pooled for further experimentation.

Next, these rtTA-expressing cell pools were infected with retroviruses carrying either pRetroX-Tight-Pur-P21-GFPcDNA or control pRetroX-TightPur. Selection of successfully transduced clones was done using puromycin, and maintenance was under 250 ng/ml of the same antibiotic. The expression of P21-GFP was monitored via Western blot and fluorescence microscopy (Bio-Rad ZOE, U.S.A.) following doxycycline administration.

### Preparation of Lentiviral Vectors for USP14, PSMD14, and Regγ shRNA

Oligonucleotides were designed for the cloning of shRNA-encoding sequences against USP14, PSMD14, and Regγ. We used USP14 shRNA primers with sense (USP14shRNA-F: GATCCCCGGCTCAGCTGTTTGCGTTGTTCAAGAGACAACG CAAACAGCTGAGCCTTTTTGGAAA) and antisense (USP14shRNA-R: AGCTTTTCCAAAAAGGCTCAGCTGTTTGCGTTGTCTCTTG AACAA CGCAAACAGCTGA GCCGGG) sequences; REGγ shRNA primers with sense (REGshRNA-F: GATCCCCGTGAGGCAGAAGACTTGGTTTCAAGAGAACCAAGTCTTTCTGCCTCACTTTTTGGAAA) and antisense (REGshRNA-R: AGCTTTTCCAAAAA GTGAGGCAGAAGACTTGGT TCTCTTGA AACCAAGTCTTCTGCCTCACGGG) sequences; and PSMD14 shRNA primers with sense (PSMD14shRNA-F: GATCCCCGTCTATATCTCTTCCCTGGTTCAAGAGACCAG GGAAGAGATATAG ACTTTTTGGAAA) and antisense (PSMD14shRNA-R: AGCTTTTCCAAAA AGTCTATATCTCTTCCCTGGTCTCTTGA ACCAGGGAAGAGATATAGACGGG) sequences. The aforementioned oligonucleotides were annealed and subcloned into the BglII-HindIII site of the pSUPER vector (VEC-PBS-0002, Oligoengine, U.S.A.). Subsequently, the BamHI-SalI fragments from these pSUPER constructs were subcloned into the same sites of pRDI292, a backbone plasmid of pLV.PARP1#5 (a gift from Didier Trono - Addgene plasmid # 14548), to produce pLV.Usp14shRNA, pLV.PSMD14shRNA, and pLV.Regγ shRNA plasmid vectors.

Hek293T cells were transfected with the shRNA plasmids (pLV.shUSP14, pLV.shPSMD14, pLV.shRegγ) along with the helper vectors psPAX2 and pMD2.G using the Calcium Phosphate Mammalian Transfection Kit (631312-Takara bio). Lentivirus-containing supernatants were collected 48 hours post-transfection and used to infect target cells in the presence of 10 mg/ml polybrene. After 24 hours, the medium was replaced with one containing 0.5-3 mg/ml puromycin for a 6-day selection period. Surviving cells were pooled and maintained with 250 ng/ml puromycin. The knockdown effects of the shRNAs were verified by Western blot analysis.

### Overexpression of USP14 using the Retro-X Tet-On Advanced Inducible Expression System

USP14 cDNA was PCR amplified from HepG2-derived cDNA using specific primers and cloned into an EcoRV-digested pBSK-clo2D vector, then excised using BamHI. The fragment was ligated into the BamHI site of the pRetroX-Tight-Pur vector for inducible expression. Construct orientation was verified before amplifying the final construct in DH5α E. Coli. The resulting virus was used to infect previously rtTA-transduced HepG2, Hep3B, and T98G cells. Selection was carried out with 0.5-3 mg/ml puromycin for 6 days, with cell maintenance in 250 ng/ml puromycin. USP14 overexpression was confirmed by Western blot.

### Preparation of Hep3B Cells Stably Expressing Ub-R-YFP

The Ub-R-YFP open reading frame was excised from an Addgene vector (#11948) using NheI and NotI, blunt-ended, and cloned into pBSK-clo2D pre-digested with EcoRV. Following cleavage with BamHI, the Ub-R-YFP insert was ligated into a similarly cut and dephosphorylated pLV-EF1a-IRES-Puro vector. Proper insertion was confirmed, and the recombinant plasmid was amplified in DH5α E. coli.

Hek293T cells were transfected with the Ub-R-YFP construct along with psPAX2 and pMD2.G using a calcium phosphate kit. Lentiviral supernatants were harvested and used to infect Hep3B cells in the presence of polybrene. Following a 24-hour infection, cells were selected using puromycin and surviving populations were pooled and maintained with puromycin.

### Preparation of Cells Stably Expressing LC3B

We generated the pLV-EF1a-IRES-Puro-LC3B vector for stable expression. LC3B cDNA was PCR-amplified from an Addgene plasmid using specific primers and cloned into EcoRV-digested pBSK-clo2D. The LC3B insert was excised with BamHI and ligated into the BamHI site of a dephosphorylated pLV-EF1a-IRES-Puro vector. After confirming the correct orientation, the construct was propagated in DH5α E. coli.

Hek293T cells were transfected with pLV-EF1a-IRES-Puro-LC3B and packaging vectors psPAX2 and pMD2.G. Lentiviral particles were harvested and used to infect target cells with the aid of polybrene. Post-infection, cells underwent selection with puromycin and maintained pools were assessed for LC3B expression via fluorescence and confocal microscopy.

### Preparation of Cells Stably Expressing APP695 Using pLV-EF1a-IRES-Puro Vector

The open-reading frame for APP695 was excised from the pc DNA FRT TO-APP695 plasmid, a kind gift from Aleksandra Radenovic (Addgene plasmid #114193), using HindIII and EcoRV. It was then cloned into pBSK-clo2D. Cloning into the pLV-EF1a-IRES-Puro vector and subsequent stable cell line preparation followed the methods previously detailed.

### Proteasome 20S Assay

We measured proteasome activity with the Enzo Life Sciences’ 20S assay kit, applying isolated and purified proteasomes from human erythrocytes.

### HIS-Tag p21 Protein Expression and Purification

The HIS tag was incorporated into p21 cDNA primers on the C-terminus. After PCR amplification, the product was cloned into the Retro-X Tet-On Advanced system and expressed in Hep3B cells equipped with the rtTA transactivator. Post-doxycycline induction, His-tagged p21 was purified using Ni-NTA Ultra-Flux Affinity Microspheres from cell lysates prepared with a Dounce Homogenizer.

### Native Gel Electrophoresis

Native gel electrophoresis was performed with protocols adapted from those described by Hoffman L. et al.**^65^**

### Brain Dissection and Sample Preparation (Digestion and Mass Spectrometry)

Brains were rapidly dissected from the subjects and immediately flash-frozen in liquid nitrogen to preserve the tissue for later analysis.

### Tissue Homogenization and Purification of Synaptosomes

We processed brains from groups of three mice, with each mouse’s synaptic fractions prepared individually. The method for synaptosome purification involved a discontinuous sucrose gradient, detailed as follows.

Brains were homogenized in HEPES-buffered sucrose with inhibitors. The homogenate was centrifuged to discard the nuclear fraction; the supernatant underwent further centrifugation to isolate the synaptosomal fraction, which was then lysed by hypoosmotic shock, homogenized, and centrifuged to remove the vesicular fraction. This preparation was used for the sucrose gradient at 4 °C.

A gradient of 0.8 M, 1.0 M, and 1.2 M sucrose in HEPES was layered and synaptosomal membranes were carefully placed on top, then centrifuged to separate the synaptic plasma membrane (SPM). The SPM was diluted, centrifuged to pellet, resuspended, and lysed in Triton X solution. After another centrifugation, the supernatant (TS fraction) and resuspended pellet were rotated, centrifuged, and the final synaptosome pellet was resuspended in HEPES/EDTA.

#### Protein Extraction from Synaptosomes

Proteins were extracted from synaptosomes using 1xRIPA buffer with inhibitor cocktails. Post-centrifugation at 32,000 x g for 30 min at 4 °C, the supernatant with the proteins was collected. Protein concentrations were determined using a BCA Assay, with BSA as a standard.

### In Solution Protein Digestion

Protein samples (100 µg each) were reduced with 5 mM DTT and alkylated with 50 mM IAA at room temperature, protected from light. After adding more DTT (to 10 mM), proteins were precipitated and redissolved in 8 M urea with 50 mM Tris.HCl, pH 8.5. Upon dilution to 1M urea with Tris.HCl, trypsin was added (1:100) and incubated overnight at 25 °C. TFA (0.5% v/v) stopped the digestion. Samples were purified, evaporated, and resuspended to 1 g/L for analysis.

#### Mass Spectrometry

The peptides underwent chromatography on a 50 cm EASY-Spray column with a 75 µm internal diameter, packed with 2 µm PepMap C18 particles (100 Å pore size). Chromatography was performed using the Ultimate 3000 RSLnano system. Analysis utilized the Q Exactive Plus mass spectrometer. Peptides were introduced onto the column with 0.1% formic acid in water (buffer A). A 150-minute gradient elution from 5% to 95% acetonitrile with 0.1% formic acid (buffer B) was carried out at a flow rate of 250 nanoliters per minute. Mass spectra were acquired using an Orbitrap Q Exactive Plus mass spectrometer in data-dependent mode with a scan range from 500 to 2500 m/z over four hours. Peptide fragmentation was achieved through Higher-energy Collisional Dissociation (HCD) at 29% normalized collision energy (NCE). MS/MS spectra were acquired at a resolution of 17,500.

### Data Analysis

MaxQuant version 1.6 facilitated concurrent protein identification and label-free quantification (LFQ). The search maintained default settings with a first search demonstrating monoisotopic mass accuracy of ±20 ppm and a second search at ±4.5 ppm. Tryptic peptides allowed up to two missed cleavages, and peptide charge was permitted to reach +7. Fixed and variable modifications included carbamidomethylation of cysteine residues (Cys), oxidation of methionine residues (Met), and N-terminal acetylation. Mus musculus protein sequences (55,087 sequences, UniProt) were used for querying raw MS data files. Perseus (v1.6) was employed to examine MaxQuant outcomes for comparability. The LFQ data of synaptosome labeled samples were combined to compare whole synaptic proteins, revealing the proteome profile of the entire synapse under Ketamine treatment.The LFQ values from MaxQuant served as primary variables for statistical analysis. The dataset was then filtered to exclude proteins solely recognized by site, potential contaminants (defined by a pre-established list), and proteins with a reversed sequence. Intensity values underwent a base 2 logarithmic transformation, and a filtering process based on valid values was applied to the rows, requiring a minimum of two valid values per row. For inclusion in the final dataset, each peptide needed to have a valid LFQ value in at least two biological replicates. Missing LFQ values were imputed by generating random numbers from a Gaussian distribution, adjusted to mimic the LFQ values of low abundance proteins. Subsequently, Perseus was used to perform a two-sample Student’s t-test between control and drug-treated mice. (Reference: Tyanova et al., 2016).

### Confocal imaging

LC was labeled with anti-LC3 antibody from Sigma Aldrich. Hoechst 33258 (Molecular Probes, Eugene, OR, USA) at a concentration of 1 mg/ml was used as a nuclear stain. Slides were examined using a Carl Zeiss laser scanning confocal microscope (LSM-510) equipped with 488-nm Argon ion, 543-nm He-Ne, and 633-nm He-Ne lasers, along with a 63× Zeiss Plan-Apo objective. The images were captured and processed using LSM-510 software from Carl Zeiss, Germany.

### Ethics Statement and Animal Experiment

The experimental protocols received ethical approval from the Ethics Committee of Ankara University Medical School Experimental Animal Care Unit (Decision Number: 2019-22-189, Date: December 11, 2019) and the Local Ethics Committee for Animal Experiments at Ankara University. The study involved adult (3-month-old) Balb/c mice, with 3 mice per group, housed at the Experimental Animal Research Laboratory of Ankara University Faculty of Medicine. All procedures were conducted in compliance with international guidelines and were approved by the Ankara University Animal Care Committee. Ketamine (Sigma-Aldrich) was administered intramuscularly at a dosage of 30 mg/kg.

### Bioinformatic Analysis

Enrichr (https://maayanlab.cloud/Enrichr/) was used for pathway analysis, integrating databases such as KEGG, REACTOME, PPI Hub proteins, and various ontology databases including GO Biological Process, Cellular Component, Molecular Function (2023 editions), MGI Mammalian Phenotype (2021 edition), and SynGO (2022 edition) for functional annotations (references 23-25). The SynGO database (https://www.syngoportal.org/) supported synaptic protein analysis (reference 21). DAVID informatics (https://david.ncifcrf.gov/) provided tools for Gene Ontology, Pathway annotations, and Functional Analysis. The String database (http://www.string-db.org/) was used for protein-protein interaction analysis (reference 41), along with the ZS Revelen-PPI tool (https://zs-revelen.com) which offered a broad scope of functional data (reference 42). FunRich (FunRic3.1.3), including a Venn diagram utility, was employed for proteomics data analysis (reference 50).

### Statistical Analysis

Data are presented as means ± standard errors of the mean (SEM). Statistical analyses were performed using Microsoft Excel (Mac v.16.59) and GraphPad Prism (v.9), employing Student’s t-test (paired, two-tailed) or Spearman rank-correlation where applicable, as specified in figure legends. Biological triplicates were standard unless otherwise noted, with the number of samples (n) indicated in figures and their legends. Results are based on a minimum of three independent experiments, with error bars representing standard deviation (SD). P-values < 0.05 were considered statistically significant, denoted as: * p<0.05, ** p<0.01, *** p<0.001, **** p<0.0001

## Supplemental information

### Legends of Figures S1 to S11

**Fig. S1** Rapid reduction of IDP-characteristic proteins by NMDAR antagonists across cell lines.

**Fig. S2** NMDAR antagonist-induced proteasome activity changes and synaptic protein alterations

**Fig. S3.** Assessment of glutamate receptors, including their antagonists and agonists on proteasome functionality and p21 levels

**Fig. S4.** The amounts of intrinsically disordered regions in p21, p53, and EYFP proteins are presented

**Fig. S5.** Exploration of the DUB enzymes USP14 and UCHL5 regarding NMDAR antagonist impact

**Fig. S6.** Evaluation of the DUB enzyme RPN11 (PSMD14) regarding NMDAR antagonist effects

**Fig. S7.** Analysis of RPN11 (PSMD14) silencing on proteasome activity in NMDAR antagonist contexts in HepG2 and T98G cell lines.

**Fig. S8.** Evaluation of H_2_O_2_ and ketamine effects on cell cycle and proteasome regulatory proteins.

**Fig. S9.** Analysis of ketamine’s effects on proteasome structures and functions.

**Fig. S10.** Characterization of protein alterations and their relation to neurodegenerative disease pathways

**Fig. S11.** Analysis of protein alterations following ketamine injection with connections to synaptic changes related to mental diseases.

### Titles of Tables S1 and S2

**Table S1:** Glutamate Receptors and Corresponding Modulating Agents Utilized in the Study

**Table S2:** Molecules screened for UPS inhibition

### Supplementary excel Tables legends

**excel Table S1**: The table reveals the altered proteins after ketamine injection, demonstrating both increased and decreased proteins that align with synapse-associated proteins outlined in the SynGO library and provided by van Oostrum *et al*.

**excel Table S2:** The table presents the cellular component information of the altered proteins, encompassing both increased and decreased proteins following ketamine injection.

**excel Table S3:** The table includes Biological Process ontology terms associated with the altered proteins, covering both increased and decreased proteins following ketamine injection.

**excel Table S4:** The table includes records from the UP_SEQ_FEATUREs chart, a subcategory of the Functional Annotation Chart from DAVID informatics sources, for altered proteins. This encompasses both increased and decreased proteins following ketamine injection, as well as randomly chosen proteins unaffected by the ketamine injection.

**excel Table S5:** The table highlights disease enrichment among all 526 altered proteins using the DisGeNET library.

**excel Table S6:** The table displays the Functional Annotation Chart of all 526 altered proteins obtained from the DAVID informatics sources.

**excel Table S7:** The table presents the HUB proteins identified using the ZS Revelen-PPI tool.

**excel Table S8**: The table unveils 231 proteins associated with diseases out of the 526 altered proteins. Furthermore, it specifies the proteins linked to schizophrenia, Alzheimer’s disease, Parkinson’s disease, and bipolar disorder within the subset of 231 disease-associated proteins.

**excel Table S9**: The table presents the HUB proteins identified from the enrichr tool.

**excel Table S10:** The table displays plasticity-related terms associated with all 526 altered proteins, obtained from various categories within the DAVID informatics sources.

